# Genome architecture and duplication-associated functional diversification in the carnivorous plant *Sarracenia purpurea*

**DOI:** 10.64898/2026.11.14.766640

**Authors:** Victor A. Albert, Jonathan Kirshner, Christopher Page, Nicholas Pratt, Johanna Merkel, Michaela Richter, Matthias Freund, Kenji Fukushima, Charlotte Lindqvist

**Author notes:** Correspondence (V.A.A.). These authors contributed equally (Jonathan Kirshner, Christopher Page).

## Abstract

The repeated evolution of complex traits raises the question of how genomes generate ecological novelty while preserving developmental organization. We present a chromosome-scale genome assembly and annotation for the carnivorous purple pitcher plant, *Sarracenia purpurea*, and investigate how two contrasting modes of gene duplication have shaped its genome and functional repertoire: ancient polyploidy, which leaves duplicated genes in conserved chromosomal regions (syntenic duplicates), and local tandem duplication, which generates neighboring gene copies. We compare these duplication classes across nine angiosperms. The genome shows a mosaic of conserved and fractionated duplicated regions reflecting differential retention and loss of genes following ancient polyploidy and later restructuring. Comparison with noncarnivorous *Actinidia* (kiwifruit) identifies 18 syntenic and nine tandem Gene Ontology (GO) Biological Process term-duplication-class combinations enriched in *Sarracenia* but not significantly enriched in *Actinidia*. Syntenic enrichments include meristem maintenance, organ- boundary specification, cuticle development, and hormonal regulation; tandem enrichments include defense responses, oxidative stress, jasmonic-acid biosynthesis, leaf senescence, and oligopeptide transport. Across the carnivorous genera *Sarracenia*, *Nepenthes*, *Utricularia*, and *Pinguicula*, we identify 19 syntenic and two tandem-associated biological processes showing significant enrichment in all four lineages, revealing striking functional commonalities across multiple evolutionary origins of carnivory despite their divergent trap morphologies and ecological strategies. Among *Sarracenia* genes differentially expressed in at least one comparison of flower, pitcher-leaf, and root tissues, tandem-class genes exhibit larger maximum absolute expression differences and greater GO richness and effective breadth at matched numbers of annotated homologs. These results highlight the complementary evolutionary contributions of ancient genome duplication and local gene-family expansion to the developmental and ecological diversification of carnivorous plants.

**Article Summary:** This study presents a chromosome-scale genome of the carnivorous pitcher plant *Sarracenia purpurea* and compares ancient genome duplication with local, tandem gene duplication. The genome contains chromosomal regions showing uneven retention and loss of duplicated genes. Compared with noncarnivorous kiwifruit (*Actinidia*), *Sarracenia* shows different patterns of enrichment for genes involved in development, defense, and stress responses. Comparisons across four carnivorous genera reveal repeated enrichment of developmental and environmental- response processes, highlighting shared functional themes across distinct evolutionary origins and trapping strategies. Among *Sarracenia* genes whose expression differs across tissues, locally duplicated genes show larger changes and a broader range of annotated biological processes.

**Graphical abstract:** 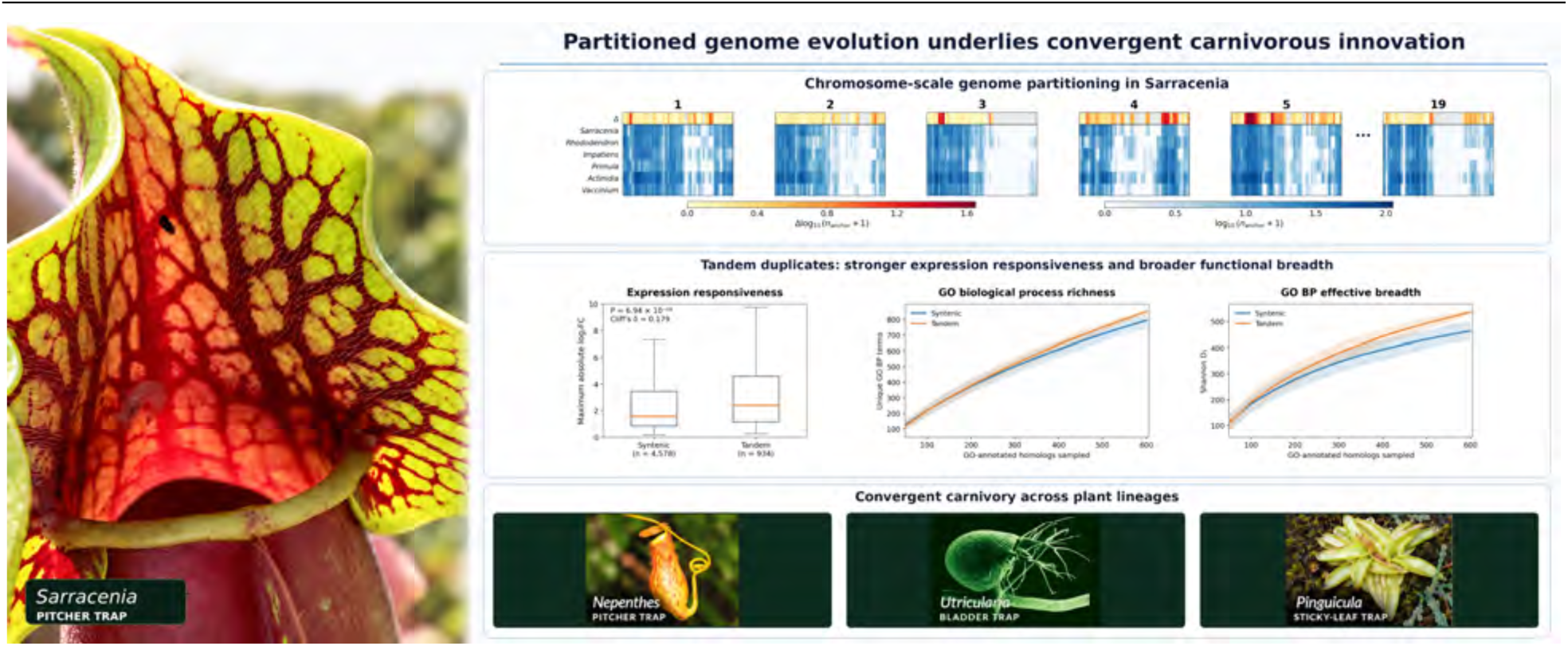

Conserved genome structure and local gene duplication contribute to distinct functional patterns in carnivorous plants. (A) Chromosome-scale comparisons show uneven retention of genes within conserved genomic regions of *Sarracenia* and related Ericales, using grapevine as a common reference. (B) Tandemly duplicated *Sarracenia* genes within the tissue-responsive subset show larger expression differences and greater diversity of annotated biological processes than genes associated with conserved syntenic regions. (C) Representative traps from *Nepenthes*, *Utricularia*, and *Pinguicula*, and comparative genomic analyses against these genera, place *Sarracenia* genomic patterns in the broader context of the repeated evolution of carnivory across flowering plants. Photos, left to right: *Sarracenia purpurea*: Victor A. Albert; *Nepenthes gracilis*: Dan Osipov (https://www.inaturalist.org/observations/354329906; CC BY 4.0); *Utricularia gibba*: Enrique Ibarra-Laclette and Victor A. Albert; *Pinguicula vulgaris*: Victor A. Albert.

## Introduction

Carnivorous plants have long attracted scientific attention because they display remarkable evolutionary innovations in leaf architecture, nutrient acquisition, and ecological strategy (Darwin 1875; Ellison and Adamec 2018). Among them, the North American pitcher plant *Sarracenia purpurea* is characterized by fluid-filled traps that capture and retain arthropod prey, whose decomposition releases nutrients available to the plant (Naczi 2018) (**Figure 1A-C**). These traps support a structured digestive ecosystem in which microbes, small invertebrates such as larval dipterans, and decomposing prey interact within a trophic network (Miner and Taylor 2002; Gray, et al. 2012; Armitage 2017; Grothjan and Young 2019; Bittleston, et al. 2020; Hale, et al. 2020; Freedman, et al. 2021; Grothjan and Young 2022; Cai, et al. 2024).

**Figure 1.**
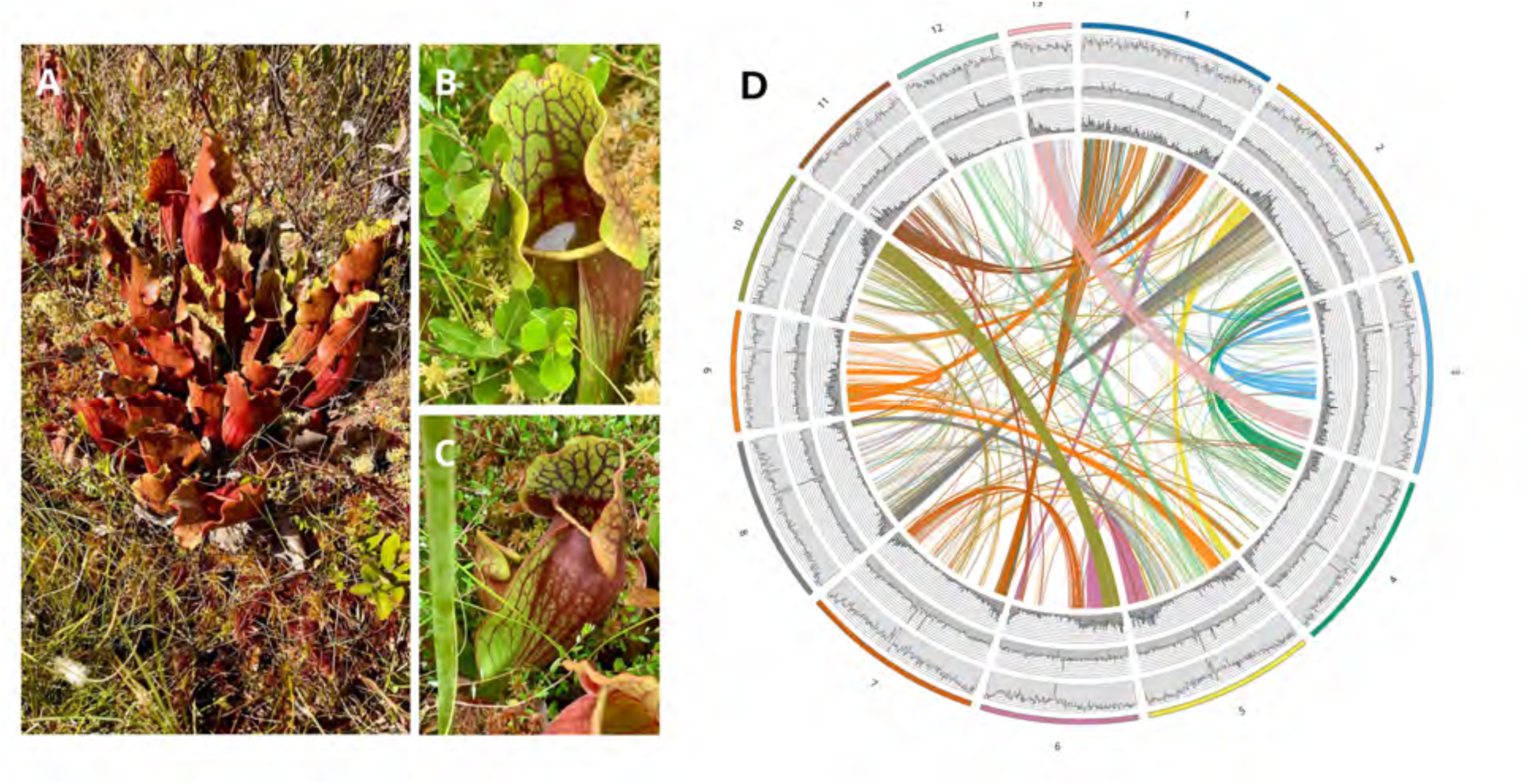
Plant habit and chromosome-wide genomic feature profiles in *Sarracenia purpurea*. (**A**) Whole plant forming a clump of pitchers, (**B**) a single pitcher showing the lid, peristome around the trap opening, and fluid-filled pitcher tube, and (**C**) a side view highlighting the conspicuous wing. (**D**) Genomic feature profiles in the *Sarracenia* genome assembly. Tracks across the 13 pseudochromosomes show - from the innermost ring outward - gene density, Gypsy transposable element (TE) abundance, and Copia TE distribution. At the center, self-syntenic blocks inferred from duplicated anchor pairs (corresponding genes retained in similar order in different genomic regions) (with log₁₀(Ks) < 1; Ks = synonymous substitution rate) are shown as color-coded chords. Photos by V.A.A.: (A), Allenburg Bog Audubon Nature Reserve, New York; (B) and (C), Mendon Ponds Park, New York.

Despite considerable phenotypic research, molecular developmental studies (Fukushima, et al. 2015), environmental research (e.g., (Segala and Horner 2023)), and marker-based phylogenetic research (e.g., (Stephens, et al. 2015; Baldwin, et al. 2023)), the genomic basis of pitcher-leaf development and digestive biology in *Sarracenia* has not been thoroughly examined. Earlier genomic investigations of carnivorous plants have addressed genome reduction and duplication in the bladderwort *Utricularia* (Lan, et al. 2017), convergent morphological and digestive adaptations in the Australian pitcher plant *Cephalotus* (Fukushima, et al. 2017), and polyploidy, sex determination, and subgenome dominance in the Southeast Asian pitcher plant *Nepenthes* (Saul, et al. 2023).

While convergence is conspicuous among pitcher plant lineages (Thorogood, et al. 2018), the genomic mechanisms by which these lineages independently reprogram leaf development and implement distinct digestive strategies remain unclear. Their contrasting digestive strategies are equally informative. For example, in *Sarracenia* (as reviewed above) nutrient release depends largely on microbial metabolism, with comparatively limited direct enzymatic contribution from the plant, whereas in *Nepenthes*, enzyme-rich secretions produced by the plant directly contribute to its carnivorous function (Kocáb, et al. 2021; Ravee, et al. 2021; Freund, et al. 2022).

We address these questions using a long-read-based, chromosome-scale assembly of *Sarracenia purpurea* and a filtered, nonredundant primary annotation in which each predicted protein-coding locus is represented by a single principal transcript. We use this resource (which was scaffolded against a recently published *Sarracenia* genome (Alabady, et al. 2026)) to assess gene retention following ancient whole-genome duplication, the gene-space contributions of local tandem duplication, and the developmental and ecological functional associations found within each duplicate gene class.

*Sarracenia* belongs to the Ericales, a diverse flowering-plant order for which an ancient allopolyploid origin - combining genetic material from divergent ancestral lineages - has been proposed (Nie, et al. 2022). The order encompasses a wide diversity of ecological strategies and well-known plants, including blueberry (*Vaccinium species*) (Rose, et al. 2018). Aligning a chromosome-scale *Sarracenia* genome with multiple Ericales relatives enables two related questions to be addressed: (1) Which duplicated blocks and structural genomic features represent the ancestral framework within which carnivory evolved? (2) Which rearrangements, patterns of duplicate-gene loss, and local gene-family expansions are lineage-specific and potentially associated with pitcher evolution?

Addressing these questions requires distinguishing gene copies associated with different duplication processes, syntenic genome comparisons, synonymous substitution rate (Ks) analysis interpreted using mixture models, and gene-level functional enrichment studies. Synteny, the conservation of gene order between chromosomal regions, identifies corresponding blocks that can be compared across species and with the ancient core-eudicot *gamma* hexaploidy event (Tang et al. 2008; Chanderbali et al. 2022). Ks measures synonymous substitutions per synonymous site between homologous coding sequences; its distribution, summarized using statistical mixture models, can provide conditional estimates of the relative timing of duplication events. Comparisons with grapevine, *Vitis vinifera*, provide a common chromosomal reference for tracking conserved gene order (Jaillon et al. 2007; Sankoff et al. 2009; Wang et al. 2025).

Within these comparisons, pairs of corresponding genes are termed syntenic anchors. Differences in the number of anchors retained in corresponding genomic regions provide a measure of local retention and fractionation, the evolutionary loss of duplicated genes. Because detectable anchor numbers can also be affected by rearrangement and sequence divergence, these patterns do not, by themselves, identify ancestral parental subgenomes.

This comparative analytical design allows tests of whether developmental regulators or stress-related gene families are preferentially represented in differently retained regions, rather than assuming such a division (Wendel et al. 2016; Bird et al. 2018; Wendel et al. 2018; Alger and Edger 2020; Twyford et al. 2025). Repeated evolution of carnivorous traps (Ellison and Adamec 2018) provides a useful comparative genomic framework. We compare *Sarracenia* with three other carnivores - *Nepenthes gracilis* (Caryophyllales), *Pinguicula gigantea* and *Utricularia gibba* (Lamiales) - and combine this sample with five noncarnivorous reference taxa, including *Arabidopsis* as the annotation reference and *Actinidia chinensis* as a noncarnivorous Ericales comparator.

For each duplication class, we record whether an identified *Arabidopsis*-mapped homolog occurs in the corresponding gene set of each of the nine sampled species. The resulting exact nine-taxon occupancy classes describe which sampled species share a homolog within that particular duplication class. Absence from an occupancy class does not necessarily indicate absence of the gene from the species’ genome.

Gene Ontology (GO) Biological Process enrichment tests then identify biological functions statistically overrepresented within these gene sets. Standardized GO enrichment was tested separately for the four carnivorous genera and *Actinidia*. *Mimulus guttatus* (monkeyflower) and *Beta vulgaris* (beet) provide noncarnivorous occupancy references for Lamiales and Caryophyllales, respectively; they were not tested as separate GO-significance filters.

The *Actinidia* comparison distinguishes functions significantly enriched in *Sarracenia* but not in the corresponding noncarnivorous Ericales gene sets. Such differences in enrichment status identify candidate functional contrasts, rather than proving that genes or biological processes are absent from *Actinidia* or originated uniquely in *Sarracenia*.

Together, we use our chromosome-scale assembly of *Sarracenia purpurea*, and its annotated coding genes and repetitive DNA, to analyze ancient duplication history represented in the *Sarracenia* genome via self-synteny, the conditional timing of inferred duplication events using synonymous substitution rates, and variation in retained syntenic gene copies using comparative anchoring against *Vitis*. Comparative tests using standardized Gene Ontology annotations examine enrichment in syntenically retained and locally tandem-duplicated gene sets, while comparative expression analyses across flower, pitcher-leaf, and root tissues quantify differences in gene expression among the sampled organs. These analyses place the carnivorous pitcher leaf within a framework of gene and genome duplication, conservation, fractionation, and tissue- associated functional diversification.

## Results and Discussion

### Chromosome-Scale Genome Assembly and Annotation

#### Genome Assembly of *Sarracenia purpurea*

The *Sarracenia purpurea* genome was assembled from high-coverage Oxford Nanopore Technology long reads (Wang, et al. 2021) and reference-scaffolded against the publicly available *Sarracenia* genome assembly of Alabady et al. (Alabady, et al. 2026) (GCA_051027295.1) to produce chromosome-scale sequences. The contig assembly spans 3,523,392,364 bp across 4,071 contigs, with a contig N50 of 2,875,369 bp. The final scaffolded assembly totals 3,523,623,004 bp and is organized into 13 chromosome-length scaffolds, or pseudochromosomes, consistent with the haploid chromosome number. Its scaffold N50 is 241.6 Mbp, and Benchmarking Universal Single-Copy Orthologs (BUSCO) analysis indicates 95.6% genome completeness and 84.1% completeness of the filtered protein annotation (**Table 1**).

**Table 1.** Comparison of genome assembly, annotation, and repeat statistics for *Sarracenia purpurea* in this study and Alabady et al. (2026). Boldface identifies selected metrics favoring the present assembly in continuity, single-copy gene representation, and reduced redundancy. Other metrics are presented descriptively. Scaffold N50 and N90 represent the minimum scaffold lengths accounting for 50% and 90% of the assembly, respectively; L50 and L90 indicate the corresponding numbers of scaffolds. Genome and predicted-protein completeness were evaluated separately using BUSCO with the eudicots_odb10 dataset (2,326 orthologs). Gene density is expressed as models per megabase pair (Mbp), average intron and exon lengths in base pairs, and repeat content as percentages of assembled sequence. The annotation in this study retains one representative transcript per protein-coding locus. Alabady et al. statistics were obtained from the published study and deposited genome and annotation (GCA_051027295.1), with comparative calculations described in Methods. Differences in annotation pipelines and transcript-model selection contribute to these contrasts, while the lower genomic duplicated-BUSCO fraction independently supports reduced assembly-level redundancy.

| METRIC | (ALABADY, ET AL. 2026) | THIS STUDY |
|---|---|---|
| **ASSEMBLY** |  |  |
| GENOME SIZE | 3.3 Gbp | 3.5 Gbp |
| NUMBER OF SCAFFOLDS | 6,330 | **1,884** |
| LARGEST SCAFFOLD | 281 Mbp | **321 Mbp** |
| SMALLEST SCAFFOLD | 1.1 kbp | **1.5 kbp** |
| SCAFFOLD N50 | 220 Mbp | **241 Mbp** |
| SCAFFOLD N90 | 0.1 Mbp | **7.6 Mbp** |
| SCAFFOLD L50 | 7 | 7 |
| SCAFFOLD L90 | 1,362 | **26** |
| **GENOMIC BUSCO (EUDICOTS)** | 2326 | 2326 |
| COMPLETE | 2134 (91.8%) | **2223 (95.6%)** |
| SINGLE | 1676 (72.1%) | **2036 (87.5%)** |
| DUPLICATED | 458 (19.7%) | **187 (8.0%)** |
| FRAGMENTED | 31 (1.3%) | **26 (1.1%)** |
| MISSING | 161 (6.9%) | **77 (3.3%)** |
| **PROTEIN BUSCO (EUDICOTS)** | 2326 | 2326 |
| COMPLETE | 92.30% | 84.10% |
| SINGLE | 72.40% | **75.80%** |
| DUPLICATED | 19.90% | **8.30%** |
| FRAGMENTED | 1.10% | 7.40% |
| MISSING | 6.60% | 8.50% |
| **ANNOTATION** |  |  |
| GENE MODEL COUNT | 52,067 | **28,835** |
| EXON COUNT | 177,368 | **132,001** |
| CDS COUNT | 172,445 | **132,001** |
| GENE DENSITY | 15.6 models/Mbp | **8.18 models/Mbp** |
| AVG INTRON NUMBER PER GENE | 2.4 | 3.6 |
| AVG INTRON LENGTH | 2221 | 2267 |
| AVG EXON NUMBER PER GENE | 3.4 | 4.6 |
| AVG EXON LENGTH | 262 | 263 |
| **REPEATS CONTENT** |  |  |
| LTR PROPORTION | 74.42% | 71.80% |
| COPIA | 29.23% | 33.40% |
| GYPSY | 23% | 20.50% |
| UNKNOWN | 25.70% | 12.40% |

Direct comparison with the previously published assembly (Alabady, et al. 2026) indicates that the present assembly provides a more continuous and less redundant representation of the genome. Although the two assemblies have similar scaffold N50 values, the present assembly requires only 26 scaffolds, rather than 1,362, to encompass 90% of the assembled sequence. It also recovers a larger proportion of complete genomic BUSCOs, 95.6% versus 91.8%, while reducing duplicated genomic BUSCOs from 19.7% to 8.0%. Notably, the Alabady et al. assembly has nearly the same duplicated-BUSCO proportion in its genomic and predicted-protein analyses, 19.7% and 19.9%, respectively. This concordance suggests that much of the excess duplication is present in the assembled genomic sequence itself and is therefore consistent with retained haplotigs or partial diploid representation, rather than arising solely from alternative splice forms in the annotation. These properties support the present assembly as a less redundant collapsed-haploid representation for structural and comparative genomic analyses. The underlying long-read contig assembly and gene annotation were generated independently. Chromosome-scale ordering and orientation, however, were guided by the Alabady et al. (Alabady, et al. 2026) assembly using RagTag, so chromosome-scale scaffold organization should not be regarded as an independent validation of the Omni-C-derived structure of that assembly.

Extended regions of reduced gene density mark putative pericentromeric and centromeric regions (**Figure 1D**), providing positional context for self-synteny and local gene-retention analysis, without determining parental subgenome identities.

#### Annotation and Gene Space Characterization

The filtered, nonredundant annotation contains 28,835 protein-coding loci, each represented by a single primary transcript model (**Table 1**). This is substantially fewer than the 52,067 gene models reported by Alabady et al. (Alabady, et al. 2026), producing gene densities of 8.18 and 15.6 models/Mbp, respectively. Some of this difference is likely attributable to the assembly-level redundancy indicated by the elevated genomic duplicated-BUSCO score in the Alabady assembly. The remaining difference may also reflect the inclusion of multiple splice forms or otherwise redundant transcript models rather than selection of one representative model per locus. Although the more inclusive Alabady et al. protein set has higher raw BUSCO completeness, 92.3% versus 84.1%, the present annotation has a higher single-copy BUSCO fraction, 75.8% versus 72.4%, and markedly fewer duplicated BUSCOs, 8.3% versus 19.9%. Thus, the present annotation is less redundant and provides a conservative estimate of the primary protein-coding gene complement. This reduced assembly and annotation redundancy is particularly important for the analyses presented below because retained haplotigs or redundant gene models could artificially inflate inferred gene-family sizes, tandem-duplicate counts, homolog occupancy patterns, and functional-enrichment signals. Transcript-supported models from trap, flower, and root tissues (see Methods, RNA-seq Processing and Expression Analyses) supported gene prediction and retained trap-associated expression contexts. Mean exon length is approximately 263 bp; genes average approximately 4-5 exons; and approximately 25% are single-exon loci (**Table S1**). Because the two studies used different annotation pipelines and transcript-model selection criteria, gene-model counts and protein BUSCO completeness should not be interpreted as directly comparable quality ranks.

Gene density is highly heterogeneous across chromosomes, with extended gene-poor regions interspersed with localized gene-rich segments (**Supplementary Figure S1**). Across the 31 analyzed scaffolds exceeding 5 Mb, 1-Mb windows contain 0 to 42 overlapping annotated genes and exhibit a strongly right-skewed distribution (**Supplementary Figure S2A**).

RepeatMasker (Chen 2004) estimates that ∼88% of the genome is repetitive, with ∼86.6% represented by interspersed repeats (**Table S2**). At the level of TE class composition, LTR (long terminal repeat) retrotransposons dominate, comprising ∼71.8% of the genome, with Copia elements representing ∼33.4% and Gypsy elements ∼20.5%. DNA transposons account for an additional ∼1.1%, while the remaining repeat fraction is distributed among unclassified elements (∼12.4%) and other low-abundance families. Together, these proportions indicate that the *Sarracenia* genome has been shaped primarily by extensive LTR retrotransposon accumulation at the genome-wide scale.

Repeat content is high across all pseudochromosomes but varies modestly among them: total TE coverage ranges from ∼84.3-90.6% (Δ ∼6.3 percentage points; coefficientt of variation ∼2%), and LTR-tagged coverage from ∼38.9-45.3% (Δ ∼6.3 percentage points; CV ∼4%). This chromosome-scale variation, together with the negative window-level association between LTR coverage and low gene density (**Supplementary Figure S2B**), is consistent with megabase-scale compartmentalization of gene-rich and repeat-rich regions.

RepeatMasker divergence landscapes summarize repeat divergence and the relative history of repeat accumulation (**Supplementary Figure S3**). Repeat fragments were binned by percent divergence from consensus (0-50%), and bp coverage was summed per bin to generate bp-weighted profiles. Genome-wide repeats show a mode near ∼7% divergence, whereas LTR-tagged repeats peak near ∼12%. The separation between genome-wide and LTR-tagged modes indicates that this explicitly LTR-tagged subset contributes to structured turnover and complement chromosome-scale tracks (**Figure 1D, Supplementary Figure S1**) by revealing heterogeneity in repeat divergence across classes.

### Internal and Comparative Syntenic Architecture

#### *Sarracenia* Self-Synteny and Structural Genomic Landscape

Self-synteny analysis using SynMap in CoGe (Lyons, et al. 2008; Haug-Baltzell, et al. 2017; Albert and Krabbenhoft 2023) reveals extensive duplicated *Sarracenia purpurea* chromosomal segments consistent with ancient polyploidy (**Figure 1D**; cf. (Srivastava, et al. 2011)). Long, nearly collinear blocks coexist with fragmented segments, consistent with gene loss, rearrangement, and TE insertion. After filtering 5,824 self-synteny Ks values to 0.05-2.0, a prespecified, biologically informed two-component Gaussian mixture (in recognition of the *gamma* hexaploidy and a *Sarracenia*-specific polyploidy) estimates means of 0.610 and 1.517 (**Figure 2**). More complex mixtures yield lower Bayesian information criterion (BIC) values and are treated as statistical decompositions rather than automatically as additional genome duplications (Lynch and Conery 2003; Cui, et al. 2006).

**Figure 2.**
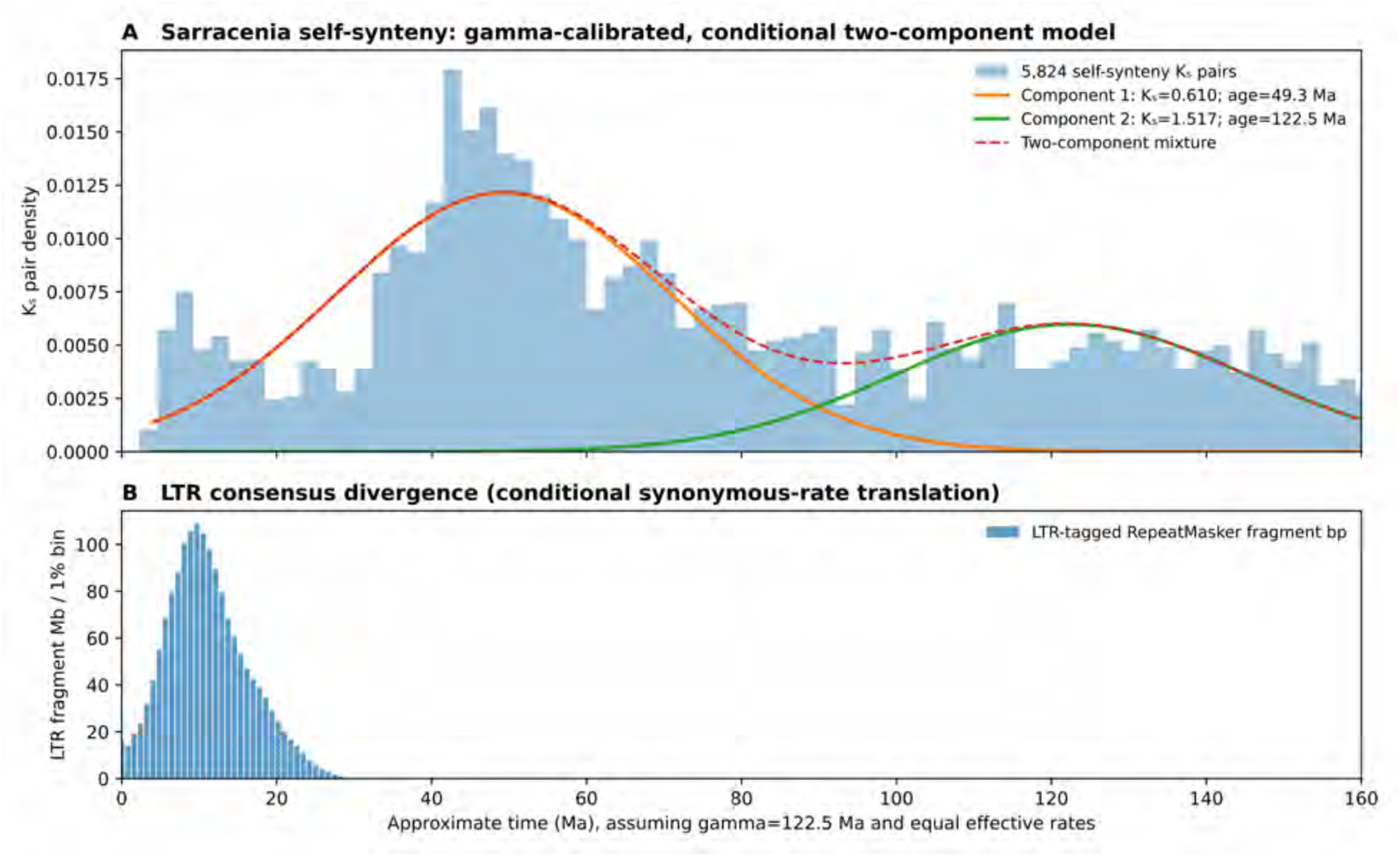
*Sarracenia* duplication and repeat-divergence signals under an explicitly specified calibration. (**A**) Distribution of 5,824 self-synteny Ks observations within 0.05-2.0, shown with a two-component Gaussian mixture (means 0.610 and 1.517) calibrated by assigning the higher-Ks component to gamma at 122.5 Ma. The lower component consequently corresponds to approximately 49.3 Ma. Two components summarize biologically interpreted broad structure but do not minimize BIC among all candidate mixture sizes. (**B**) RepeatMasker LTR-tagged fragment bases by percent consensus divergence, translated to an illustrative age axis using the same effective substitution rate. The LTR divergence peak is 12%; its inferred approximately 9.7 Ma timing remains rate- dependent.

Assigning the older of these two Ks components to the core-eudicot gamma calibration of 122.5 Ma gives a conditional age estimate of 49.3 Ma for the younger component (**Figure 2**). The LTR-tagged RepeatMasker fragment distribution peaks at 12% consensus divergence; applying the effective substitution rate inferred from the Ks calibration yields an illustrative age of approximately 9.7 Ma for the LTR peak. Because synonymous substitutions and repeat divergence need not evolve at identical rates, this is a relative comparison rather than an independently dated insertion event. Both timing estimates depend on model choice and calibration (Jiao, et al. 2012; Vekemans, et al. 2012; Wang, et al. 2016).

#### Multispecies Ericales Comparative Anchoring and Duplication Timing

To evaluate duplication timing in a broader context, Ks distributions were examined across Ericales taxa. All sampled species retain a broad, high-Ks component consistent with the core eudicot *gamma* hexaploidy. Some taxa, including *Actinidia* and *Vaccinium*, also exhibit prominent lower-Ks components (results not shown), likely reflecting more recent duplication events. Because these younger events obscure separation between *gamma*-derived and post-*gamma* signals that may be Ericales-general, *Actinidia* and *Vaccinium* were not used for cross-taxon comparison, although these genomes remain structurally informative.

Across *Sarracenia*, *Rhododendron*, *Primula*, and *Impatiens*, self-synteny Ks landscapes contain broad components that can be summarized by two-component mixtures, although model complexity and component assignment remain uncertain (**Supplementary Figure S4**). Using 0.05-2.0 Ks and independently constraining the older component to gamma at 122.5 Ma yields younger conditional estimates near 49.3 Ma for *Sarracenia*, 64.2 Ma for *Rhododendron*, 73.3 Ma for *Primula*, and 62.3 Ma for *Impatiens*. Alabady et al. (Alabady, et al. 2026) reported approximately 81-84 Ma for *Sarracenia* paleopolyploidization. Their duplicate selection, decomposition, and calibration differ, so these estimates are not directly comparable without standardized reanalysis.

**A**lthough the ages for post-*gamma* events detected in *Rhododendron* and *Impatiens* are similar, differences in peak breadth, prominence, and modeling sensitivity among all taxa prevent confident assignment of homology among events. These patterns reveal substantial post-gamma duplication activity across Ericales, although the available Ks distributions do not resolve whether the younger components derive from shared or lineage-specific events.

#### Syntenic Architecture and Local Retention Relative to *Vitis*

Comparative synteny across Ericales shows extensive correspondence with the *Vitis vinifera* outgroup reference. All six Ericales comparisons were projected onto the same *Vitis* CoGe genome 19990 coordinates. Mapping *Sarracenia* onto *Vitis* recovers large intervals of syntenic correspondence with local differences in retained anchor density, compatible with ancient polyploidy, fractionation, and subsequent rearrangement (Joyce, et al. 2017; Albert and Krabbenhoft 2023). Across 19 *Vitis* reference chromosomes, each subdivided into 50 bins, the *Sarracenia* structural analysis identifies 688 bins with a unique highest-retention chromosome track, 55 ties at the highest rank, and 207 bins without qualifying finite-Ks chromosome anchors (**Figure 3**). Comparisons of six Ericales taxa on the common *Vitis* reference demonstrate conserved and lineage-variable anchor patterns (**Figure 3**).

**Figure 3.**
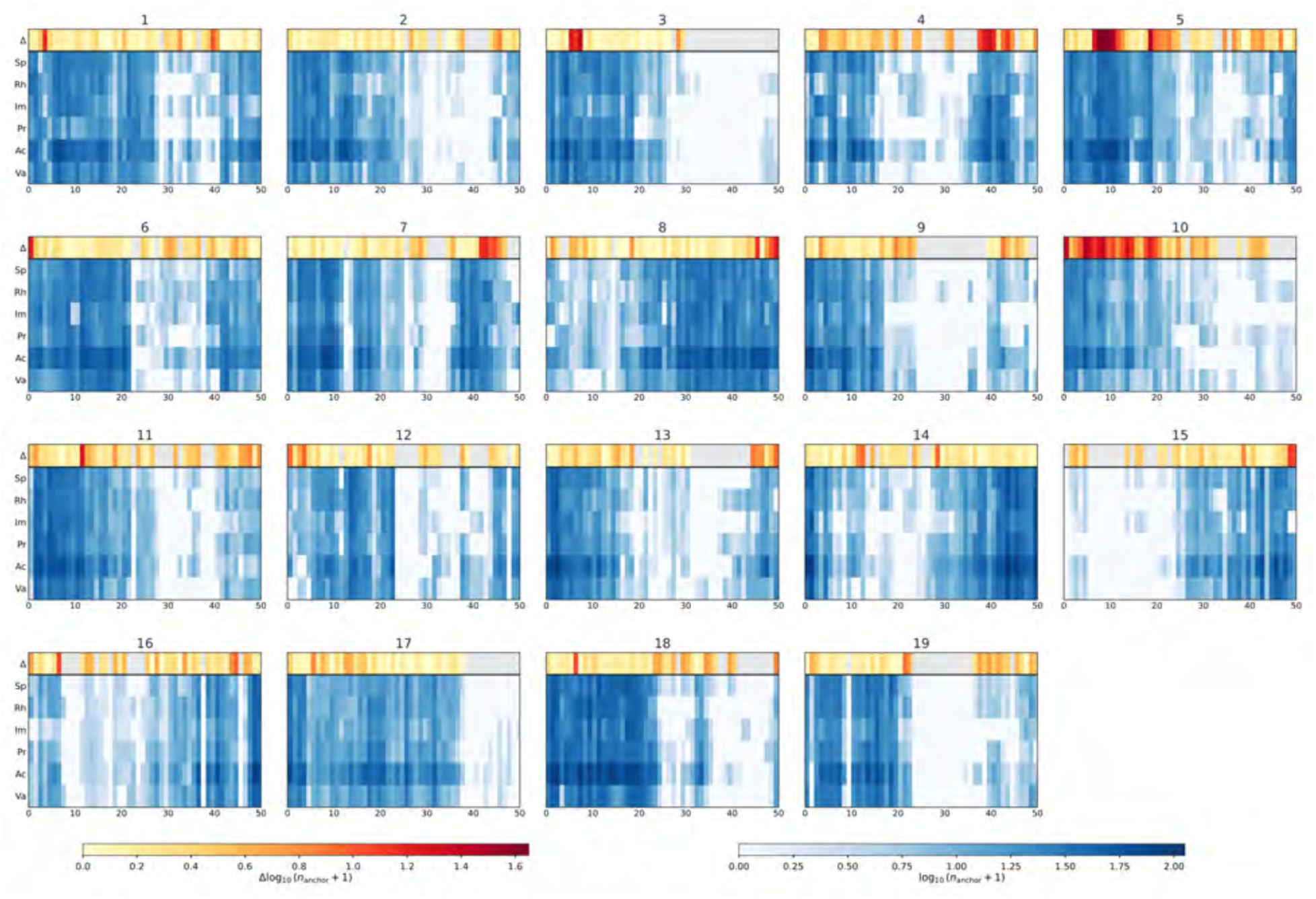
Chromosome-scale patterns of syntenic anchor retention and local genomic asymmetry across six plant lineages. Each of the 19 panels represents a chromosome of the *Vitis vinifera* reference genome (CoGe genome ID 19990), divided into 50 equal physical-coordinate intervals. Horizontal axes indicate relative bin position (0-50), rather than physical distance in megabases or gene number. The upper, yellow-to-red track (Δ) represents the local difference in log-transformed syntenic anchor counts between the highest-ranking and second-highest-ranking *Sarracenia* chromosomes within each reference interval: Δ = log₁₀(*n*ₘₐₓ + 1) − log₁₀(*n*__second_+ 1). Larger values (orange to red) indicate greater disparity in local anchor retention. Zero indicates equal highest counts, whereas gray denotes intervals lacking qualifying *Sarracenia* anchors. The six lower tracks show syntenic anchor densities for Sp, *Sarracenia*; Rh, *Rhododendron*; Im, *Impatiens*; Pr, *Primula*; Ac, *Actinidia*; and Va, *Vaccinium*. Blue intensity represents log₁₀(*n*ₐₙ_c_ₕₒᵣ + 1), where *n*ₐₙ_c_ₕₒᵣ is the number of syntenic anchor pairs assigned to the corresponding reference interval. Darker blue indicates higher anchor counts, while white or very pale blue indicates zero or low counts. The warm-colored Δ track and blue comparative tracks use separate quantitative color scales, shown below the panels; the blue scale is shared across all six taxa. These distributions describe regional differences in syntenic anchor retention and local genomic asymmetry but do not establish the parental identities of ancestral subgenomes.

Local syntenic-retention rankings are not ancestral parental-subgenome assignments. Of the 743 occupied *Vitis* bins, 234 have qualifying anchors on only one *Sarracenia* chromosome; these yield the maximum top-versus-mean-of-other-tracks score even though no second positive track is represented. Accordingly, we report local top-versus-second contrasts and distinguish tied or unanchored bins, rather than identifying lower-ranked intervals as genetically recessive ancestral compartments. Independent genome-wide windows show a negative association between self- syntenic anchor density and repeat coverage (Spearman’s ρ ≈ −0.47 for total repeat coverage).

Fixed-bin profiles show a mosaic of higher and lower *Sarracenia* anchor retention across the *Vitis* reference (**Supplementary Figures S5 and S6**). This pattern is consistent with heterogeneous fractionation and local rearrangement, but its assignment to distinct progenitor-derived subgenomes would require longer-range collinear block phasing and a statistically supported parental-origin model. The current analyses do not provide that phase.

### Comparative Functional Architecture of Carnivory

#### Functional Architecture from Homolog Occupancy Classes Across Nine Taxa

To compare duplicate-gene repertoires across carnivorous and noncarnivorous relatives, we constructed a nine-taxon framework comprising the carnivorous taxa *Sarracenia*, *Nepenthes*, *Pinguicula*, and *Utricularia* and the noncarnivorous reference taxa *Actinidia*, *Vitis*, *Arabidopsis*, *Mimulus*, and *Beta*. *Actinidia* provides the sampled noncarnivorous Ericales comparison (Fleck and Jobson 2023). *Mimulus* and *Beta* provide noncarnivorous Lamiales and Caryophyllales references (Fleck and Jobson 2023) for duplicate-homolog occupancy, not separate tests of GO enrichment. *Vitis* anchors deeper structure, and *Arabidopsis thaliana* supplies standardized homolog identifiers and GO annotation.

For each duplication mode, Arabidopsis homolog identifiers were placed into exact nine-taxon occupancy categories describing their presence within the available syntenic or tandem duplicate datasets. Membership in such a category reflects the sampled duplicate-class data; absence from a category does not establish whole-genome absence. Species-level foregrounds are defined as unions of mutually exclusive occupancy categories.

We assigned Arabidopsis homolog IDs to genes in the syntenic and tandem sets, and encoded occupancy separately for the nine sampled taxa. Overlap categories and enrichment tests were computed on these defined gene sets rather than treating duplicate-class absence as genome-wide gene loss.

We used a fixed Arabidopsis Gene Ontology annotation framework for genome-derived enrichment. For each carnivorous taxon and each duplication class, GO Biological Process terms were tested against the same annotated background, and Benjamini-Hochberg adjustment was performed over the same predefined set of GO terms. Comparisons across taxa were then made among terms independently significant within each taxon; a union signifies enrichment in at least one lineage, whereas an intersection requires enrichment in every named lineage (**Figure 4**).

**Figure 4.**
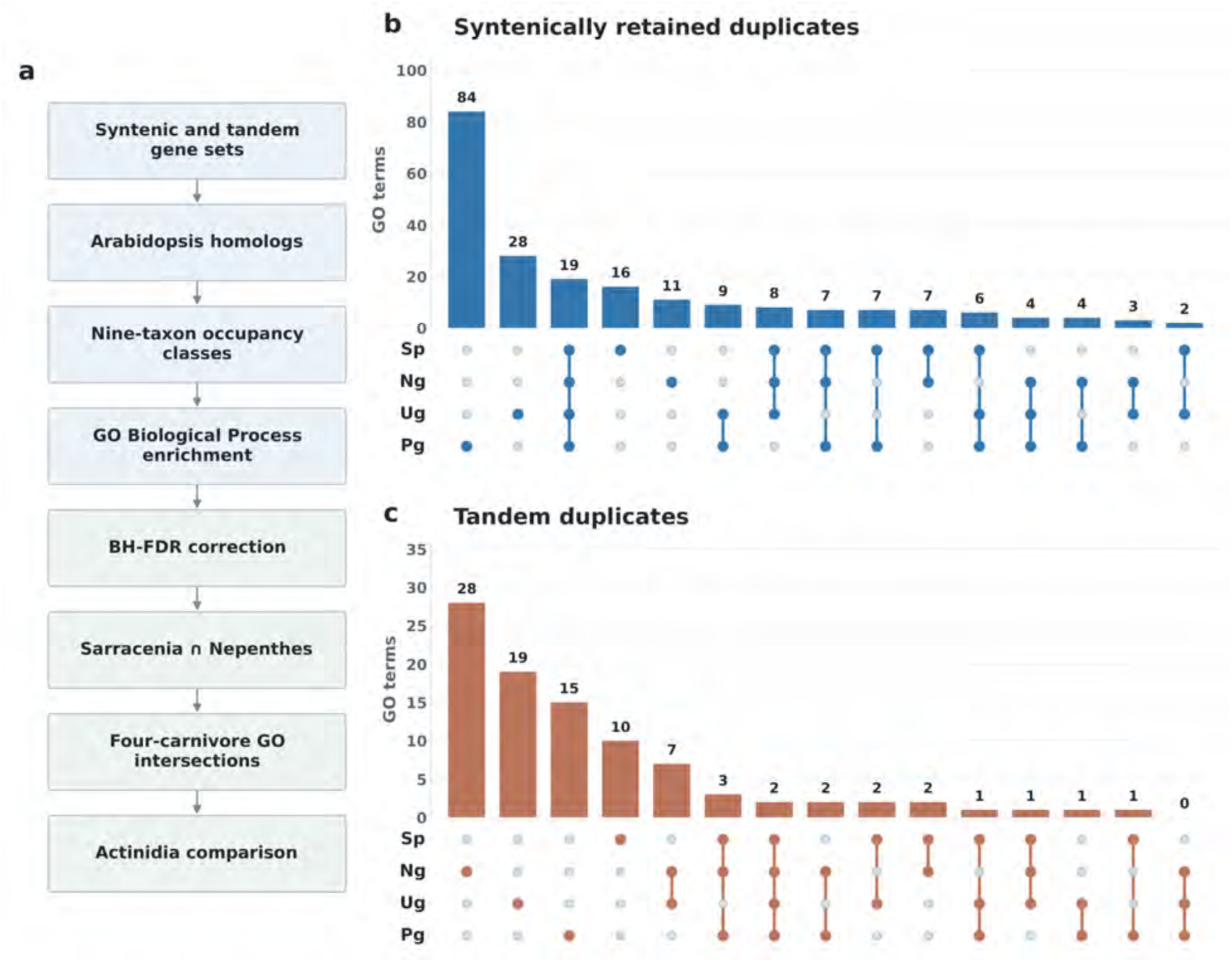
Standardized comparative enrichment of GO Biological Process terms in carnivorous angiosperms. (**A**) For syntenic and tandem duplication sets, Arabidopsis homologs were assigned to exact nine-taxon duplicate-class occupancy patterns. Each species and duplication class was tested with a one-sided hypergeometric overrepresentation test against 23,167 Arabidopsis genes annotated to at least one eligible Biological Process term. A fixed family of 3,708 active directly annotated terms was evaluated per species and duplication class; Benjamini-Hochberg correction was applied independently to each family and terms with q < 0.05 were retained. (**B**) Syntenic and (C) tandem GO terms significant in the four carnivorous genera *Sarracenia* (Sp), *Nepenthes* (Ng), *Utricularia* (Ug) and *Pinguicula* (Pg). Bars and the membership matrix show all 15 mutually exclusive patterns of significance, including 19 shared syntenic and two shared tandem terms. Counts refer to separately significant GO terms, not raw GO annotation occurrences. Expression-derived GO analysis is independent of these genome-derived tests.

The *Actinidia* comparison was applied after within-taxon testing: categories significant in *Sarracenia* but not significant in the corresponding *Actinidia* duplicate class were tabulated separately. Lack of significance in *Actinidia* must not be equated with biological absence or a uniquely derived *Sarracenia* process.

These comparisons reveal recurring functional enrichments across independently evolved carnivorous lineages, while distinguishing shared biological processes from the particular genes contributing to their enrichment.

#### Functional Enrichment Contrasts Between *Sarracenia* and *Actinidia*

The standardized GO Biological Process analysis identifies 18 syntenic and nine tandem term-duplication-mode combinations significantly enriched in *Sarracenia* (BH q < 0.05) but not significantly enriched in *Actinidia* for the same duplication mode (**Table S3**). The syntenic set includes meristem maintenance, organ-boundary specification, vegetative phase change, cuticle development, abscisic acid signaling, and vascular patterning. Tandem enrichments instead include defense and oxidative-stress responses, glutathione metabolism, jasmonic-acid biosynthesis, leaf senescence, and oligopeptide transport. The annotations derive from Arabidopsis homologs; representative *Sarracenia* gene models are catalogued in **Tables S8 and S9**.

These 27 term-mode associations reflect differences in the outcomes of two independently performed within-taxon enrichment tests: each meets q < 0.05 in *Sarracenia* but q ≥ 0.05 in *Actinidia*. These 27 term–mode associations distinguish the enrichment profiles of the sampled duplication classes, rather than establishing the evolutionary absence or origin of individual biological processes (**Table S3**).

Across the four carnivorous genera, GO enrichments show both overlap and considerable lineage-specific differences in sampled duplicate-class functional repertoires. These patterns support recurrent use of broad developmental and environmental response processes while leaving gene-level mechanisms to be resolved by comparative functional analyses.

#### Shared Functional Enrichments in *Sarracenia* and *Nepenthes*

The *Sarracenia*-*Nepenthes* comparison tests whether two independently evolved pitcher-trap lineages show enrichment of similar Gene Ontology Biological Process categories. Each species and duplication class was tested separately against the same 3,708-term annotated background, then significant term sets were compared by intersection (enrichment in both taxa) and union (enrichment in either taxon) (**Table S5**). Of 153 term/mode entries significant in at least one lineage, 41 syntenic and eight tandem categories are significant in both. Shared associations include aspects of leaf development, stress signaling and biosynthetic regulation; the overlap identifies shared functional themes across independent pitcher lineages.

*Sarracenia* and *Nepenthes* differ in digestive strategy, with the former relying more heavily on microbial mediation and the latter producing substantial secreted hydrolytic activity. Despite their contrasting digestive strategies, both pitcher genera exhibit significant enrichment of developmental and environmental-response processes, revealing functional commonalities across independently evolved pitfall traps.

#### Conserved Duplicate-Associated Functions Across Four Carnivorous Genera

Across the four carnivorous genera examined - *Sarracenia*, *Nepenthes*, *Pinguicula*, and *Utricularia* - 19 syntenic and two tandem GO Biological Process terms meet the significance criterion independently in every genus (**Figure 4**; **Table S6**). These four genera span three evolutionary origins of carnivory because *Pinguicula* and *Utricularia* are both Lentibulariaceae. The same 21 term-mode combinations are also significant in the noncarnivorous comparator *Actinidia*; they therefore highlight broadly conserved functional repertoires potentially available for recruitment during carnivory evolution. The larger union of terms significant in any single carnivorous genus must not be interpreted as a four-way intersection.

The four genera also differ in which processes reach the specified enrichment threshold within syntenic and tandem sets (**Tables S5-S9**). Together, these patterns reveal a substantial shared functional repertoire across multiple origins of carnivory, alongside lineage-specific differences in the biological processes associated with syntenic and tandem duplication.

#### Interpretation of the *Sarracenia*-*Actinidia* Enrichment Contrast

*Actinidia chinensis* provides an ericalean noncarnivorous comparison for within-taxon enrichment tests, while *Mimulus guttatus* and *Beta vulgaris* were included in the broader homolog-occupancy framework rather than the standardized GO testing families. The *Sarracenia*-*Actinidia* comparison is a comparison of statistical enrichment status, not a direct test of whether a function evolved in either species.

For this comparison, 18 syntenic and nine tandem Biological Process term-mode entries meet the *Sarracenia*-significant/*Actinidia*-nonsignificant criterion (**Table S3**). Unlike the four-carnivore shared GO terms that are also enriched in *Actinidia*, these filtered terms highlight development and epidermal structure among syntenic sets and defense, stress, and metabolism among tandem sets. These contrasts nominate different development-related and environmental-response categories for functional follow-up.

The *Sarracenia*–*Actinidia* comparison thus highlights contrasting functional enrichment profiles within syntenic and tandem gene sets, identifying developmental, epidermal, defense, and stress-related processes as prominent components of *Sarracenia* genome evolution. (**Table S3**).

### Regulatory Deployment in Pitcher Tissue

We analyzed gene expression in *Sarracenia* separately to assess the tissue-associated deployment of functions associated with different modes of gene duplication. Differentially expressed genes (DEGs) were identified between flowers and pitcher leaves, mapped to *Arabidopsis* homologs using the same orthology framework, and subjected to GO enrichment. Because differential expression reflects regulatory deployment rather than taxonomic occupancy, presence-absence occupancy classes were not applied; instead, enriched GO terms from DEG sets were compared qualitatively with occupancy-class modules.

Transcriptome comparisons identify several homologs represented in the nine-taxon occupancy classes are also strongly expressed in pitcher leaves. Within the full DEG set, multiple *Sarracenia* homologs of *PEN3*, *ZIP6* (Robe, et al. 2025), and *DML1* display significant trap-biased expression (negative flower vs. leaf (FvL) log₂FC), consistent with xenobiotic export, micronutrient transport, and redox-responsive chromatin regulation (**Tables S11 and S12**). Several pan-carnivore homologs (AT1G01580, AT1G24140, AT2G16390) also show trap-biased expression for at least one paralog (**Tables S11, S12, and S6**). The heatmap and volcano plots (**Figure 5; Supplementary Figure S7**) highlight the strongest FvL contrasts, while genes such as *PEN3*, *ZIP6*, and *DML1* exhibit consistent trap-biased expression across the broader DEG distribution. Flower-biased expression includes terpene synthases (*TPS03* (Michael, et al. 2020), *TPS-CIN* (Roos, et al. 2015)), peroxidases (*PER7* (Vijayakumar, et al. 2016)), lipid-transfer proteins (*LTP4* (Brotman, et al. 2012)), cytochrome P450s (*CYP76G1* (Kondou, et al. 2008), *CYP71A25 (Wu*(Wu, et al. 2025)), and canonical floral organ identity regulators (*PI*, *SEP1*, *SEP4*, *AGL16* (Nam, et al. 2003; Par̆enicová, et al. 2003)) (**Tables S11 and S12**). The flower-biased expression of these floral regulatory homologs, together with the contrasting pitcher-biased expression of transport, stress-response, and metabolic genes, highlights the distinct transcriptional programs deployed in reproductive and carnivorous leaf tissues.

**Figure 5.**
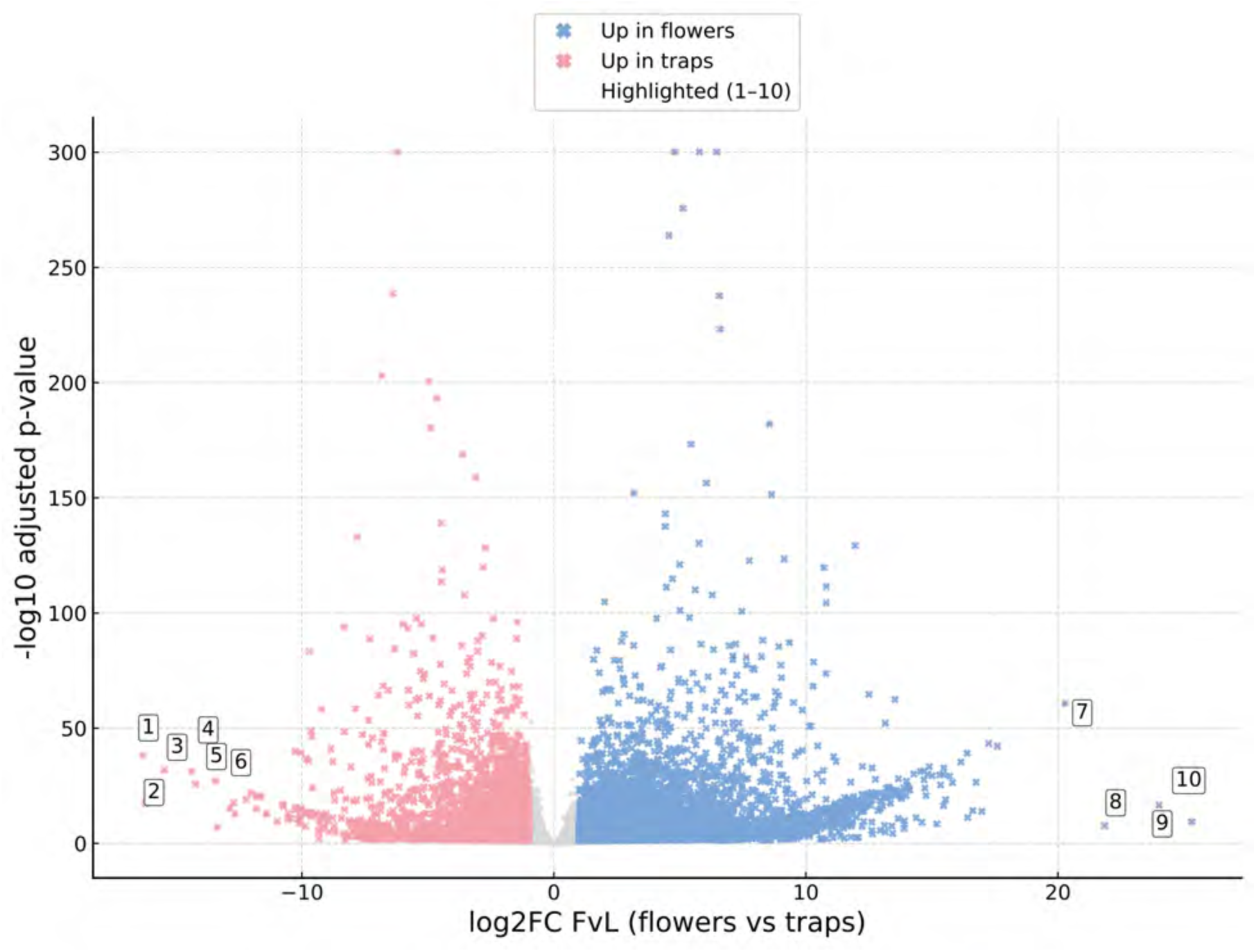
Differential expression between flowers and pitcher leaves in *Sarracenia purpurea*. Volcano plot showing differential expression between flowers and pitcher leaves. Positive log₂ fold changes (log₂FC) indicate higher expression in flowers, whereas negative values indicate higher expression in pitcher traps. The vertical axis represents −log₁₀(FDR-adjusted P value). Blue points represent flower-upregulated genes (n = 5,403), pink points represent trap-upregulated genes (n = 3,302), and gray points represent genes not meeting the significance criteria (n = 9,024). For this plot, colored upregulation classes required FDR < 0.05 and |log₂FC| > 1; the underlying DEG significance threshold was adjusted P < 0.05. Four adjusted P values recorded as zero were assigned a plotting value of 300 on the −log₁₀ scale. The six genes with the largest negative log₂FC values and the four with the largest positive values are numbered 1-10, respectively. Functional descriptions are based on their best *Arabidopsis thaliana* sequence matches and do not imply experimentally established functions in *Sarracenia*. 1. **AT3G23230 (*TDR1*)** — trap-biased ERF transcription factor. 2. **AT1G04220 (*KCS2*)** — trap-biased 3-ketoacyl-CoA synthase. 3. **AT3G19820 (*DWF1*)** — trap-biased brassinosteroid biosynthesis-associated protein. 4. **AT1G53540 (*HSP17.6C*)** — trap-biased small heat-shock protein. 5. **AT1G70170 (*MMP*)** — trap-biased matrix metalloproteinase. 6. **AT3G09110** — trap-biased protein of unknown function. 7. **AT5G59310 (*LTP4*)** — flower-biased lipid-transfer protein. 8. **AT1G30870 (*PER7*)** — flower-biased peroxidase. 9. **AT3G25830 (*TPS-CIN*)** — flower-biased terpene synthase. 10. **AT4G16740 (*TPS03*)** — flower-biased terpene synthase.

Trap-biased transcripts reveal gene-expression patterns associated with the physiological specialization of pitcher leaves. Strongly trap-biased transcripts include *TDR1* (Zhang, et al. 2012), *KCS2* (Lian, et al. 2021), *DWF1* (Best, et al. 2016), *HSP17.6C* (Yamaguchi, et al. 2021), *MMP* (Wu, et al. 2022), reflecting cuticle biosynthesis, brassinosteroid metabolism, stress buffering, osmotic regulation, and proteolytic remodeling. Additional regulators such as *CER1* (Yang, et al. 2020), *CBF4* (Guttikonda, et al. 2014), *WRKY9* (Krishnamurthy, et al. 2021), *PCNA1* (Strzalka, et al. 2015), *PRL* (Holding and Springer 2002), and *DAO1* (Zhang, et al. 2016) support integration of surface modification, stress-response, hormonal, and redox modules.

Flower-enriched tandem-only homologs (AT1G02205, AT1G55260, AT1G62045) indicate tissue-biased expression of defense-associated homologs in reproductive tissues (**Tables S11 and S12**). Distinct paralogs of *PEN3* and *ZIP6* occur in both trap- and flower-biased sets, suggesting divergence in tissue-associated expression among these duplicated transporters.

Together, these transcriptional profiles reveal tissue-specific deployment of genes represented in the comparative duplication datasets. The contrasting expression of genes associated with transport, defense, hormone signaling, metabolism, and surface formation highlights the functional specialization of pitcher leaves and the integration of developmental and physiological processes underlying carnivory.

### Integrated Model for *Sarracenia* Pitcher Evolution

*Sarracenia* retains numerous homologs of established developmental regulators, including AGO1 (Bohmert, et al. 1998), BRX (Beuchat, et al. 2010), GATA11 (Manfield, et al. 2007), ETC1 (Tominaga-Wada, et al. 2017), and RCD1 (Shapiguzov, et al. 2019), within conserved syntenic regions (**Table S12**). Their Arabidopsis annotations are consistent with retention of important developmental functions after ancient duplication, but do not demonstrate ancestral parental-subgenome identity or preferential dosage-regulator retention. In a gene-level comparison of 436 matched *Vitis* reference bins, six GO-based functional categories concerning developmental regulation, transcription factors, stress response, defense, proteolysis, and biotic response showed no significant enrichment between highest- and second-ranked syntenic tracks after BH correction (all q > 0.59).

The tandem gene sets include loci annotated with detoxification, glutathione-mediated redox, defense, cuticle and cell-wall processes (**Tables S4, S8 and S9**). Such annotations support potential ecological functions, but the present tests do not demonstrate that these genes accumulate preferentially in lower-retention or parentally recessive segments.

Among *Sarracenia* genes significantly differentially expressed in at least one flower, pitcher-leaf, or root contrast, the duplication-class dataset contains 934 tandem and 4,578 syntenic genes. Within this tissue-responsive subset, responsiveness was defined for each gene as the maximum absolute log₂ fold change across the flower-versus-leaf, root-versus-leaf, and flower-versus-root contrasts. Tandem genes exhibited a higher median responsiveness than syntenic genes (2.410 versus 1.568; two-sided Mann-Whitney P = 6.94 × 10⁻¹⁸; Cliff’s δ = 0.179; **Figure 6A**).

**Figure 6.**
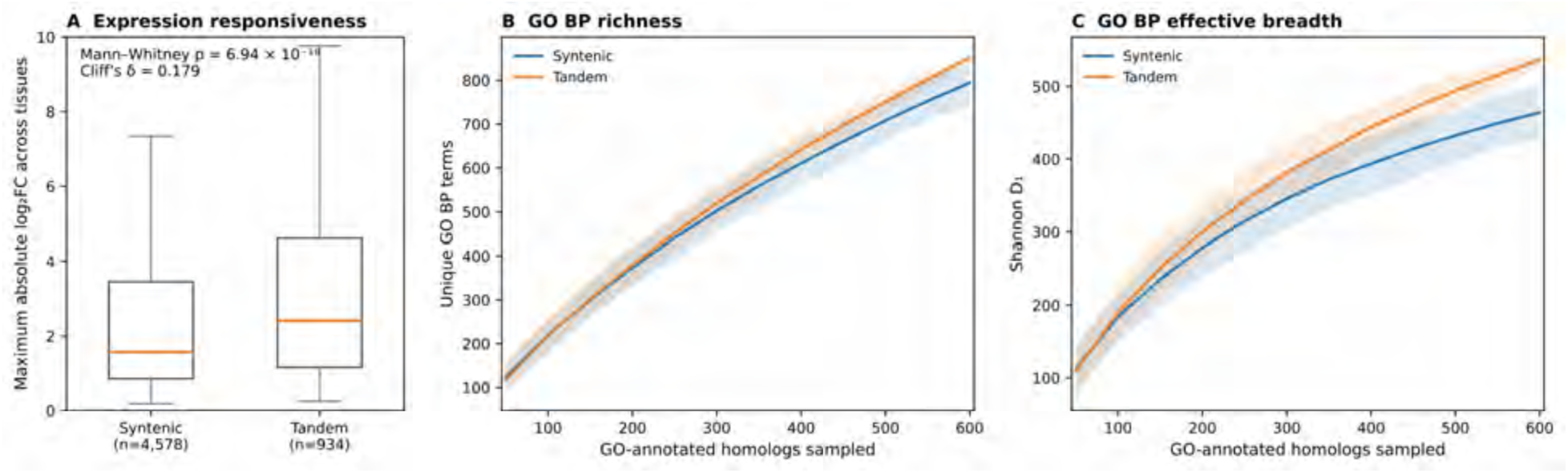
Expression responsiveness and GO Biological Process breadth among tissue-responsive tandem and syntenic *Sarracenia* gene sets. (A) For 934 tandem and 4,578 syntenic genes with significant adjusted P values in at least one of three pairwise tissue contrasts, responsiveness is the largest absolute log2 fold change among flower versus leaf, root versus leaf, and flower versus root. The two-sided Mann-Whitney P is 6.94 × 10^-18 (Cliff’s delta = 0.179). Boxes show medians and interquartile ranges; whiskers exclude plotted outliers. (B) Mean distinct GO Biological Process richness and (C) Shannon effective breadth (D1) among 617 tandem-associated and 2,305 syntenic-associated GO-annotated Arabidopsis homologs. At each depth, 300 random without-replacement subsamples were drawn per class; curves show archived means and shaded 2.5th-97.5th percentile bands among subsamples. These bands describe resampling variability and are not independent biological confidence intervals. The rarefaction panels describe functional annotation diversity within the selected tissue-responsive subset; no significance test using the dependent resamples is reported.

Rarefaction was used to compare functional annotation diversity between the expression-responsive *Sarracenia* duplication classes at equivalent sampling depths. Repeated subsampling of distinct Arabidopsis homolog identifiers minimized differences arising from unequal gene-set sizes and annotation redundancy. Direct GO Biological Process annotations were available for 617 tandem-associated and 2,305 syntenic-associated homologs. At an equal sample size of 500 annotated homologs, tandem sets had greater mean GO richness (751.0 versus 708.1 terms) and Shannon effective breadth (the exponential of Shannon entropy, D₁) (494.1 versus 432.4; **Figure 6B,C**). Thus, tandem-associated genes exhibit both greater functional annotation richness and greater functional breadth than syntenic genes within the tissue-responsive subset, even after standardizing the number of annotated homologs.

*Sarracenia* and *Nepenthes* share 41 syntenic and eight tandem GO-term/mode associations independently significant within both lineages. The broader union of terms significant in either species comprises 153 mode/term entries (**Table S5**). This substantial overlap reveals shared developmental, regulatory, and environmental-response functions across two independently evolved pitcher-plant lineages, despite their contrasting strategies for prey digestion and nutrient acquisition.

Under the gamma-based calibration, the younger *Sarracenia* Ks component corresponds to approximately 49.3 Ma, whereas the principal LTR divergence peak corresponds to approximately 9.7 Ma on the same calibrated time scale. This pronounced temporal separation supports a genome evolutionary history in which ancient gene duplication was followed by a substantially more recent phase of LTR retrotransposon accumulation.

Together, the analyses reveal complementary dimensions of duplicated-gene evolution in *Sarracenia*: a chromosome-scale mosaic of syntenic retention and fractionation, enrichment of developmental and structural processes in syntenic duplicate sets relative to the noncarnivorous *Actinidia* comparison, and enrichment of defense- and stress-related categories among tandem duplicates. Within the expression-responsive gene subset, tandem duplicates show greater tissue-associated expression changes and a richer, more even GO Biological Process repertoire at matched annotated-homolog sample sizes (**Figure 6; Table S3**). The four-carnivore GO comparison additionally identifies recurring duplicate-associated functional enrichments across multiple origins of carnivory (**Figure 4**). These complementary structural, comparative, and expression results support a model in which inherited genomic organization and local duplicate diversification both contribute to the evolutionary potential for novel leaf functions.

### Comparative Implications and Evolutionary Synthesis

The chromosome-scale assembly of *Sarracenia purpurea* reveals how ancient genome duplication, differential gene retention, and local tandem duplication have shaped the genomic repertoire of a carnivorous plant. Extensive syntenic correspondence with other Ericales lineages provides a conserved chromosomal framework, while self-synteny and a younger post-*gamma* duplication signal support an additional ancient polyploidy event in the lineage leading to *Sarracenia*, potentially shared more broadly across Ericales. Heterogeneous gene retention within duplicated chromosomal regions reveals extensive subsequent fractionation and evolutionary restructuring.

The extraordinary abundance of LTR retrotransposons in *Sarracenia*, accounting for approximately 72% of the assembled genome, reveals a second major dimension of genome evolution beyond ancient polyploidy. The separation between the younger gene-duplication signal and the principal LTR divergence peak under the illustrative shared-rate calibration suggests distinct episodes of genome duplication and subsequent repeat accumulation. Together with the concentration of repeats in gene-poor regions, these patterns indicate that retrotransposon proliferation contributed substantially to the expansion and structural compartmentalization of the modern *Sarracenia* genome.

The functional consequences of these different duplication histories are reflected in the contrasting biological processes associated with syntenic and tandem gene sets. The *Sarracenia– Actinidia* comparison identifies 18 syntenic and nine tandem GO Biological Process term– duplication-mode combinations significantly enriched in *Sarracenia* but not in the corresponding *Actinidia* sets (**Table S3**). These contrasts highlight meristem and organ-boundary regulation, cuticle development, and hormonal signaling among syntenic genes, and defense, oxidative-stress responses, glutathione metabolism, and jasmonate biosynthesis among tandem genes. Together, these patterns reveal contrasting functional associations of the two duplication modes in *Sarracenia*, emphasizing developmental and structural processes among syntenic duplicates and environmental-response and metabolic processes among tandem duplicates.

Broader comparisons place these functional associations within the repeated evolution of carnivory across flowering plants. Across *Sarracenia*, *Nepenthes*, *Pinguicula*, and *Utricularia*, 19 syntenic and two tandem GO Biological Process term–mode combinations are significantly enriched in all four genera, representing three independent evolutionary origins of carnivory (**Figure 4; Tables S5–S9**). The enrichment of these same processes in noncarnivorous *Actinidia* identifies a broadly conserved functional repertoire that may have provided evolutionary opportunities for carnivorous specialization. More specifically, the independently evolved pitcher plants *Sarracenia* and *Nepenthes* share 41 syntenic and eight tandem enrichments, revealing substantial functional commonalities despite their contrasting digestive strategies. These comparisons highlight both shared developmental and environmental-response functions and lineage-specific elaboration of the gene repertoires associated with different trapping strategies.

Tissue-specific expression provides an additional dimension of functional diversification within this genomic framework. RNA-seq comparisons among flowers, pitcher leaves, and roots identify tissue-biased *Sarracenia* genes, including pitcher-biased homologs of *PEN3*, *ZIP6*, and *DML1*, together with genes associated with environmental responses, nutrient transport, and surface modification. These expression patterns connect the comparative duplication analyses with the distinctive physiology of carnivorous leaves. Among 934 tandem and 4,578 syntenic genes in the expression-responsive subset, tandem duplicates exhibit larger maximum absolute tissue contrasts (**Figure 6A**), complemented by greater GO richness and effective breadth when annotated homologs are sampled at equal depth (**Figure 6B,C**). Thus, tandem-associated genes exhibit not only distinctive functional enrichments but also greater tissue-dependent transcriptional responsiveness and a broader annotated functional repertoire within the expression-responsive subset.

The evolutionary significance of these patterns can be investigated further through comparisons with additional independently evolved pitcher plants. A future chromosome-scale genome for *Cephalotus* would extend chromosome-level comparisons to a third independently evolved pitcher-trap lineage, allowing more detailed examination of shared gene-family histories, conserved developmental pathways, and lineage-specific genomic changes. Together with comparative resources from other carnivorous plants, such analyses would help distinguish the evolutionary contributions of ancestral functional repertoires from subsequent gene-family diversification and regulatory specialization.

Taken together, these findings reveal complementary dimensions of genome evolution in *Sarracenia*. Ancient polyploidy established a duplicated chromosomal framework subsequently reshaped by fractionation and extensive retrotransposon accumulation, while local tandem duplication contributed gene families enriched for defense, environmental responses, and metabolism. The stronger tissue-associated expression changes and greater functional annotation breadth of tandem genes add a regulatory dimension to this structural contrast. Rather than a strict separation of developmental and ecological functions into distinct genomic compartments, the resulting picture is one of conserved genomic organization combined with diversification of gene families and their expression. This interplay between inherited genome structure, local duplication, and regulatory specialization provides an evolutionary framework for understanding the emergence of novel leaf functions in carnivorous plants.

## Materials and Methods

### Genome Resources and Analytical Scope

All analyses used the chromosome-scale genome assembly of *Sarracenia purpurea* comprising 13 pseudochromosomes (https://genomevolution.org/coge/GenomeInfo.pl?gid=69349) and its corresponding representative protein-coding gene annotation. For gene-density profiling, a gene was counted in every nonoverlapping 1-Mb genomic window with which its annotated interval overlapped, rather than assigned only by its midpoint. Repeat features were used as interval annotations for chromosome- and window-based summary statistics. The RepeatMasker source assembly comprised 2,042 sequences totaling 3,523,999,544 bp, compared with 1,884 scaffolds totaling 3,523,623,004 bp in the final assembly. Repeat-class percentages and nonoverlapping masked-base coverage are reported relative to the RepeatMasker input length (3,523,999,544 bp), rather than the final assembly length.

### Genome Construction and Annotation Workflow

#### Plant Material, DNA Extraction, and Oxford Nanopore Sequencing

Plant material and voucher specimens for sequencing were collected from mature individuals of *Sarracenia purpurea* at Mendon Ponds Park, Rochester, New York (**Figure 1**). Species identification was performed in the field. Voucher specimens are currently deposited at the University at Buffalo. Leaf tissue was placed into individual vials and stored at −80 °C until DNA extraction.

Approximately 10 g of flash-frozen tissue was used for isolation of high-molecular-weight (HMW) genomic DNA. Nuclei isolation followed the BioNano NIBuffer protocol, in which frozen leaf tissue was homogenized in liquid nitrogen and subjected to a nuclei lysis step using IBTB buffer supplemented with spermine and spermidine and filtered immediately prior to use. IBTB buffer consisted of Isolation Buffer (IB; 15 mM Tris, 10 mM EDTA, 130 mM KCl, 20 mM NaCl, 8% (m/V) PVP-10, pH 9.4) containing 0.1% Triton X-100 and 7.5% (v/v) β-mercaptoethanol (BME), mixed and chilled on ice. The homogenized tissue-buffer mixture was strained to remove undissolved plant material. Triton X-100 was then added to a final concentration of 1% to lyse nuclei, followed by centrifugation at 2000 × g for 10 min to pellet the nuclei. Following nuclei isolation, DNA was extracted using a cetyltrimethylammonium bromide (CTAB) protocol modified for Oxford Nanopore sequencing. The quality and concentration of HMW genomic DNA were assessed using a NanoDrop spectrophotometer and agarose gel electrophoresis following standard protocols. Extracted genomic DNA was further purified using a Qiagen Genomic-Tip 500/G according to the manufacturer’s instructions. Aliquots of purified DNA were subjected to treatment with Short Read Eliminator (SRE; PacBio) to perform size selection and enrich for DNA fragments longer than 10 kb.

High-molecular-weight genomic DNA was thereafter sequenced using Oxford Nanopore Technology (ONT) platforms. Library preparation and sequencing followed standard ONT protocols appropriate for high-molecular-weight plant genomic DNA. Sequencing generated long-read data without pre-filtering for read length or read quality. All genome assemblies and downstream analyses were performed using unfiltered ONT reads, unless explicitly stated otherwise, in order to preserve information from repeat-rich and structurally complex regions of the genome.

#### Genome Assembly

Unfiltered Oxford Nanopore reads were assembled using hifiasm v0.25.0 with the --ont flag enabled, which activates the assembler’s long-read mode optimized for nanopore data. Assembly was performed to generate a collapsed primary (haploid) assembly, rather than phased haplotypes, reflecting the study’s focus on genome structure, gene content, and duplication history rather than allelic variation.

Assembly completeness was evaluated using BUSCO v5.4.3 with the *eudicots_odb10* dataset, which assesses the presence and copy number of conserved single-copy orthologs expected in eudicot genomes. Assembly contiguity and size statistics were evaluated using QUAST v5.2.0.

To assess read support and facilitate downstream curation, unfiltered ONT reads were aligned back to the initial assembly using minimap2 v2.24, employing presets appropriate for long-read alignment. Resulting alignments were sorted using SAMtools v1.16.1.

#### Contamination Screening and Assembly Curation

The initial genome assembly was screened for potential contamination using BlobTools v1.1.1 in conjunction with BLAST+ v2.12.0. For each contig, taxonomic assignments were inferred from BLASTN similarity searches. Contigs classified as Streptophyta were retained, as were contigs lacking confident taxonomic assignment (no-hit or undefined). Contigs assigned to non-plant taxa were removed from further analyses.

Unfiltered ONT reads aligning to the retained contigs were extracted and reassembled using hifiasm with the same parameters as the initial assembly. This produced a curated assembly in which non-plant sequences were excluded while preserving read depth and long-range genomic structure across retained regions. Assembly quality and completeness were reassessed using BUSCO and QUAST as described above.

#### Homology-Based Scaffolding

The curated assembly was scaffolded using RagTag v2.1.0, employing a published and publicly available *Sarracenia purpurea* genome assembly (NCBI accession GCA_051027295.1) (Alabady, et al. 2026) as a reference guide for contig ordering and orientation. RagTag was used strictly in reference-guided scaffolding mode; nucleotide sequences were not modified, and no gap filling or sequence correction was performed.

This scaffolding step produced a chromosome-scale assembly comprising 13 pseudochromosomes and 1,871 unplaced scaffolds. Downstream genome-coordinate analyses used this RagTag-scaffolded assembly.

Because the Alabady et al. assembly was used solely as an ordering and orientation guide, comparisons of chromosome-scale scaffold organization are not fully independent tests of chromosomal structure. By contrast, assembly length, BUSCO copy state, unplaced-scaffold representation, and all annotation statistics derive from the independently assembled sequence and independently generated annotation reported here.

#### Genome Annotation

Repetitive elements were identified using a combination of RepeatModeler v2.0.4 and EDTA v2.0.1, capturing both de novo repeat families and structurally defined transposable element classes. Repeat libraries generated by the two approaches were merged and collapsed into a non- redundant repeat library using CD-HIT v4.8.1, applying thresholds of 95% sequence identity and 90% alignment coverage. The merged classified custom library (clustered_lib.fa) was used with RepeatMasker v4.1.5 and RMBlast v2.14.0+. Original RepeatMasker alignment (.out), summary (.tbl), and coordinate-level GFF results are included in the computational reproduction archive. Protein-coding gene prediction was performed using GeMoMa v1.9.0 with default parameters. Reference proteomes from five angiosperm species were used to guide prediction: *Arabidopsis thaliana*, *Camptotheca acuminata*, *Actinidia eriantha*, *Camellia lanceoleosa*, and *Vaccinium darrowii*. These species were selected to provide phylogenetic breadth within eudicots while remaining relevant to Ericales. RNA-seq data were incorporated as evidence for the prediction of splice sites.

This annotation was refined using EVidenceModeler (EVM) v2.1.0 to standardize naming and retain the best-supported primary gene model at each locus. The resulting EVM-integrated GFF3 annotation served as the final gene annotation used for all downstream analyses. Annotation completeness was assessed using BUSCO v5.5.0 on the predicted protein set.

### Syntenic Anchor Processing and Coordinate Binning

#### Syntenic Anchor Preprocessing

Syntenic anchor tables were generated in CoGe SynMap (Lyons, et al. 2008; Haug-Baltzell, et al. 2017; Albert and Krabbenhoft 2023) with FractBias (Joyce, et al. 2017) enabled. Source Ks-bearing anchor files and FractBias gene/sliding-window reports were retained. Multispecies coordinate analyses use Vitis vinifera CoGe genome 19990 as the common reference for all six Ericales taxa; the Vaccinium darrowii query uses genome 64402. FractBias reports were used for cross-checks, while the fixed 50-bin matrices were reconstructed from coordinate-bearing anchors.

- DAGChainer: relative gene order = nucleotide distance; maximum distance between two matches = 20 genes; minimum number of aligned pairs = 5 genes
- Skip random/unknown chromosomes
- Syntenic depth: Quota Align (Tang, et al. 2011)
- Ratio of coverage depth: *Vitis* = 1 against all other taxa = 4
- Overlap distance: 40
- Window size: 100 genes; use all genes in target genome

Each anchor record obtained from SynMap (at the link: “Results with synonymous/non-synonymous rate values”) contained identifiers for the source and target chromosomes, genomic coordinates for both genes, and a Ks value. FractBias results were downloaded at the “Fractionation Bias synteny report” and “Fractionation Bias sliding window results” links.

Anchors were filtered as follows:

- Anchors lacking defined chromosomal coordinates were excluded.
- Anchors mapping to unplaced or unanchored scaffolds were excluded.
- Anchors missing a Ks estimate were omitted from the primary *Sarracenia* local-retention score and all Ks mixture fitting, but retained for descriptive chromosome correspondence. The structural six-species *Vitis* reference comparison includes chromosome-anchored records whether or not Ks is available; the sensitivity of local-retention ranks to including records with missing Ks estimates was assessed separately.

#### Reference-Coordinate Binning of Syntenic Anchors

All reference-based analyses were conducted in a fixed bin coordinate system.

This reference-coordinate framework was applied uniformly wherever syntenic anchor positions were summarized along the *Vitis* genome, including both lineage-specific and multispecies analyses, where anchors were interpreted descriptively as indicators of syntenic presence rather than as direct measures of retained gene dosage.

For each reference chromosome *c*:

Let *L_c_* be the maximum coordinate observed among all anchors on chromosome *c*.

Define a fixed number of bins *B*, held constant across all chromosomes for a given analysis.

Define the bin width:

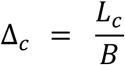

For each anchor with midpoint coordinate *m* on chromosome *c*, assign the anchor to bin:

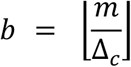

Clip *b* such that:

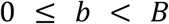

This produces a discrete bin index for every anchor, enabling direct comparison across chromosomes and taxa.

#### Anchor Density Aggregation and Transformation

For each *Sarracenia* chromosome *i* and reference bin *j*, the raw anchor count is:

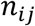

To stabilize variance and reduce domination by anchor-rich bins, counts were transformed as:

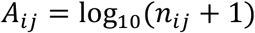

**Local syntenic retention and descriptive fractionation contrasts were calculated** from log10(anchor count + 1) values for the 13 *Sarracenia* pseudochromosomes in each Vitis bin. A ranked track is only a locally higher-or lower-retention correspondence, not an assignment to an ancestral parental subgenome. Multispecies comparative matrices are descriptive anchor-presence or density summaries.*A_i_*_j_

### Syntenic Retention Metrics and Summary Statistics

Local Syntenic Retention Margin and Fractionation Contrast

For each Vitis reference bin, the 13 *Sarracenia* chromosome-specific log-transformed anchor signals were ranked. A strictly positive difference between the highest and second-highest values indicates a uniquely highest-retention chromosome track; ties and bins with zero anchors were coded separately.*j*

Let:

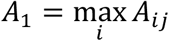

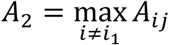

Where:

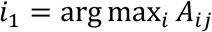

then the local top-versus-second retention margin is:

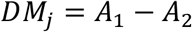

Bins with at least one anchor were classified by whether the highest value was unique or tied. Bins without qualifying anchors were coded as unanchored rather than recessive.

- A unique top-ranked chromosome track has a positive difference between the largest and second-largest log-transformed count. This is not parental subgenome dominance.*DM*_j_ > 0
- A zero top-versus-second margin denotes a tied maximum when anchors exist, or no information when every track is zero. Neither state is assigned to a recessive parental subgenome.*DM*_j_ ≤ 0

No parental-origin or dominance state was inferred from these ranked scores.

A second descriptive retention-contrast score (cf. (Joyce, et al. 2017)) was calculated as the contrast between the highest log-transformed count and the mean of the remaining 12 chromosome tracks:

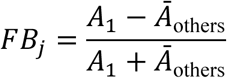

Here the comparison mean includes zero-valued tracks; values approaching 1 are therefore partly sensitive to bins containing anchors on only one *Sarracenia* chromosome. A separate top-versus-second contrast was plotted in **Figure 3**. Neither score is used to phase ancestral subgenomes. *Ā*_others_*A_i_*_j_ ≥ 00 ≤ *FB*_j_ < 1 *A*_1_ ≥ *Ā*_others_

#### Syntenic Retention Similarity Matrices

Each *Sarracenia* chromosome was represented as a vector:

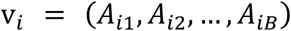

Pairwise similarity was computed using correlation:

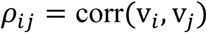

Bins with missing values were excluded pairwise. Similarities were assembled into square matrices and visualized as heatmaps.

### Ks-Based Duplication Analyses

#### Ks Filtering and Mixture Decomposition

Ks values associated with syntenic anchors were filtered to retain:

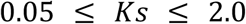

Filtered Ks values were modeled using Gaussian mixture models (Schlueter, et al. 2004; Cui, et al. 2006) fitted by expectation-maximization.

Given Ks values {*x*_1_, …, *x_n_*}, the model is:

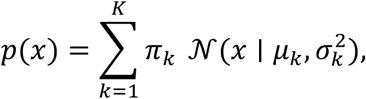

where *π_k_* denotes the mixture weight and *μ_k_* and *σ_k_* represent the mean and standard deviation of component *k*. Mixture fitting was performed using expectation-maximization, and model complexity was chosen conservatively to capture interpretable duplication signal rather than maximize statistical fit.

Model selection was performed using Bayesian Information Criterion:

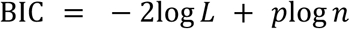

where *p* = number of estimated parameters (for 1D GMM with *k* components):

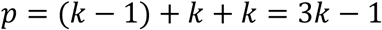

Components were sorted by increasing *μ_k_* and treated strictly as relative age classes.

#### Event-Stratified Synteny Density

Each anchor was assigned to a Ks component.

For each component *e*, reference chromosome *c*, and bin *b*:

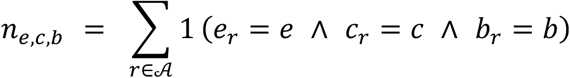

where:

• *r* indexes syntenic anchor pairs

• A is the set of all anchors retained for analysis

• *k_r_* is the Ks mixture component (event) assigned to anchor *r*

• *c_r_* is the reference chromosome to which anchor *r*maps

• *b_r_* is the reference-coordinate bin containing anchor *r*

• 1(⋅) is the indicator function (1 if true, 0 otherwise)

The per-chromosome probability normalization, i.e. each chromosome’s event-stratified density integrates to 1 across bins, is written as:

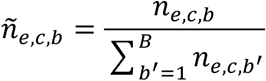

where:

• *n_e_*_,*c*,*b*_ is the raw event-stratified anchor count (as previously defined)

• ñ*_e_*_,*c*,*b*_ is the normalized density

• *B* is the total number of bins on chromosome *c*

• the denominator sums only across bins on the same chromosome and event

As such, event-stratified anchor counts were normalized within each reference chromosome so that the summed density across bins equals one. These normalized densities were used for plotting age-stratified retention profiles.

#### Ks Analysis and Conservative Duplication Inference Across Ericales

Ks distributions were analyzed to characterize the temporal structure of gene duplication within *Sarracenia purpurea* and to place these patterns within a comparative Ericales context. Duplicate gene pairs were obtained from self:self CoGe SynMap analyses(Lyons, et al. 2008; Haug- Baltzell, et al. 2017; Albert and Krabbenhoft 2023) for each species. Only gene pairs with defined chromosomal coordinates and valid Ks estimates were retained.

Ks values were filtered as above (see **Ks Filtering and Mixture Decomposition)** to reduce the influence of very recent noise at low divergence and extensive saturation at high divergence. Ks values outside this range were retained for visualization where informative but excluded from statistical modeling when dominated by multiple-hit effects. Filtered Ks values for each species were modeled using Gaussian mixture models, as in that same section above.

For all Ericales examined, the oldest and broadest Ks component was interpreted as retention of deeply diverged paralogs associated with the ancient core-eudicot *gamma* hexaploidy. This component was identified independently for each species and treated as a shared ancestral reference rather than evidence for recent duplication. Additional components at lower Ks were considered candidate post-*gamma* duplication signals only when they were distinct from the *gamma* component and not dominated by saturation or diffuse background signal.

Several Ericales taxa, including *Actinidia* and *Vaccinium*, exhibit complex Ks profiles with prominent mid-to low-Ks peaks consistent with additional lineage-specific whole-genome duplications reported for these genera. The presence of these recent WGDs obscures separation between *gamma*-derived and post-*gamma* duplication signal and complicates direct cross-taxon comparison of younger Ks modes. Accordingly, *Actinidia* and *Vaccinium* were retained for syntenic anchoring and contextual visualization but excluded from comparative inference of post-*gamma* duplication timing.

Comparative timing analyses therefore focused on *Sarracenia purpurea*, *Rhododendron*, and *Impatiens*, which lack strong evidence for additional recent polyploidy and exhibit simpler Ks landscapes. For these taxa, a single younger Ks component was modeled in addition to the *gamma* component. Component means were treated strictly as relative temporal markers rather than discrete event boundaries.

To place Ks components on a comparative time axis, Ks values were converted to divergence time using:

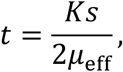

where *μ*_eff_ is an effective long-term synonymous substitution rate. For each species, *μ*_eff_ was calibrated independently by constraining the mean of the *gamma* component to 120-125 million years ago, corresponding to widely inferred estimates for the timing of core-eudicot *gamma* hexaploidy (Jiao, et al. 2012; Vekemans, et al. 2012; Wang, et al. 2016). Younger component ages were then scaled relative to this species-specific calibration.

Because Ks-based age estimates are sensitive to mixture model behavior, saturation effects, and the quality of genome assemblies and annotations, inferred duplication times are reported as approximate and are interpreted comparatively rather than as precise chronological estimates. No assumption of homology among younger duplication components across taxa was imposed.

### Repeat Analyses

#### Repeat Divergence Landscape Analysis

Repeat divergence landscapes were generated from RepeatMasker annotations to provide a relative temporal summary of repeat accumulation in the *Sarracenia purpurea* genome. All RepeatMasker-annotated repeat fragments were treated as independent genomic intervals and analyzed without reference to gene models, syntenic structure, or chromosome identity.

For each annotated repeat fragment *r*, RepeatMasker reports a percent sequence divergence from the corresponding consensus element, reflecting accumulated substitutions since insertion relative to the library consensus. Let *d_r_* denote this reported divergence value (in percent).

Repeat fragments were binned into integer divergence classes spanning 0-50% divergence. Formally, each fragment *r* was assigned to divergence bin:

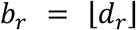

such that:

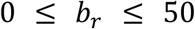

For each divergence bin *b*, total base-pair coverage was computed as the sum of fragment lengths:

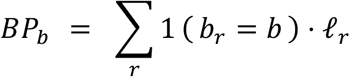

where ℓ*_r_*is the genomic length of fragment *r*. These base-pair totals were then normalized by the total assembled genome size *G* to obtain bp-weighted divergence profiles:

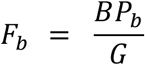

This procedure yields a genome-wide repeat divergence landscape in which each bin represents the fraction of the genome occupied by repeats with divergence values falling in that interval.

In addition to the genome-wide landscape, an LTR-restricted divergence landscape was constructed by applying the same binning and aggregation procedure to the subset of RepeatMasker fragments whose repeat names explicitly contain “LTR.” This name-tagged subset is narrower than the complete group classified as LTR retrotransposons in the RepeatMasker .out and .tbl files. All normalization and bin definitions were identical between genome-wide and LTR-restricted analyses, enabling direct comparison of their divergence profiles.

For visualization, divergence landscapes were plotted as histograms of *F_b_* across divergence bins, with smoothed curves overlaid to highlight modal structure. The smoothed curves were used solely for visualization and did not alter the underlying bin counts. The modal divergence bin for each landscape was identified as the bin *b* with maximal *F_b_* and indicated graphically as a reference.

Because RepeatMasker divergence values reflect sequence divergence from consensus rather than absolute insertion times, divergence landscapes were interpreted as relative summaries of repeat accumulation dynamics. Therefore, divergence differences involving the LTR-tagged subset describe that selected subset rather than the divergence landscape of all RepeatMasker-classified LTR retrotransposons.

#### Comparative Analysis of Ks-Based Duplication Signals and LTR Divergence Landscapes

To examine the temporal relationship between gene duplication and transposable-element accumulation within the *Sarracenia purpurea* genome, we integrated Ks-based duplication age estimates with RepeatMasker-derived divergence landscapes for the explicitly LTR-tagged subset. Ks values were obtained from self-syntenic duplicate gene pairs, and in this case, filtered to the range 0.05 ≤ *Ks* ≤ 2.0 to minimize the influence of saturation at high divergence.

Filtered Ks values were modeled using a two-component Gaussian mixture model, representing a younger lineage-specific duplication signal and a broader, older component corresponding to deeply diverged paralogs. The latter component is interpreted as reflecting retention from the ancient core-eudicot *gamma* hexaploidy event, while the younger component captures post-*gamma* duplication retained within the *Sarracenia* lineage. Mixture fitting was performed as in section 4.5.1.

The LTR-tagged divergence landscape was constructed from RepeatMasker fragments whose repeat names explicitly contain “LTR,” rather than every fragment classified as an LTR retrotransposon. For each fragment, the reported percent divergence from its consensus sequence was binned into integer divergence classes, and base-pair coverage was summed per bin to generate bp-weighted divergence profiles.

To enable direct visual comparison between duplication and repeat accumulation, Ks values and LTR divergence bins were transformed to a common time axis using the relationship:

Where:

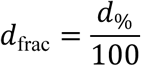

and then:

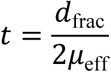

and *d* denotes Ks or fractional divergence (see also Ks Analysis and Conservative Duplication Inference Across Ericales, above). An effective long-term substitution rate *μ*_eff_ was calibrated by constraining the mean of the older Ks component to 120-125 million years ago, corresponding to the widely inferred timing of the core-eudicot *gamma* hexaploidy event. All time estimates are therefore calibration-dependent and interpreted comparatively. Ks density curves and LTR divergence histograms were plotted on aligned time axes with independent y-axis normalization to emphasize relative temporal structure rather than absolute magnitude.

#### Quantification of Transposable-Element Heterogeneity Across Pseudochromosomes

To quantify variation in transposable-element (TE) distribution across the *Sarracenia purpurea* genome, RepeatMasker annotations were summarized at the pseudochromosome scale using union base-pair coverage. All RepeatMasker-annotated intervals were parsed by pseudochromosome and merged prior to summation to avoid double-counting overlapping fragments. Total TE coverage was calculated as the number of base pairs covered by any RepeatMasker annotation (all TE classes combined) divided by the total length of each pseudochromosome.

LTR-tagged repeat coverage was calculated analogously by restricting annotations to fragments whose repeat names contain “LTR,” including tagged Gypsy and Copia entries, and computing union base-pair coverage relative to pseudochromosome length. Thus, heterogeneity was quantified separately for (i) overall TE occupancy and (ii) LTR-specific occupancy, enabling direct comparison between genome-wide repeat content and the dominant repeat class.

Inter-pseudochromosome variation was summarized using descriptive statistics across the 13 pseudochromosomes. For each metric, the range (Δ) was defined as the difference between the maximum and minimum coverage values, expressed in percentage points. The coefficient of variation (CV) was calculated as the standard deviation divided by the mean coverage and is reported as a percentage, providing a normalized measure of heterogeneity that accounts for differences in mean repeat abundance. These statistics were used to support statements regarding scaffold-level variation in repeat content and to contextualize the spatial distribution of LTR retrotransposons relative to overall TE composition.

Fragment counts per pseudochromosome were recorded as a measure of annotation density but were not used for coverage estimation. All reported percentages reflect union coverage and therefore represent conservative estimates of repeat occupancy.

### Comparative Functional Genomics (Occupancy Classes, Foregrounds, GOs)

#### Homolog Occupancy Class Construction Using Strict Intersection Logic

Protein-coding genes were first mapped to *Arabidopsis thaliana* homolog identifiers, providing a common reference framework for cross-taxon comparison. Homolog occupancy classes were then constructed using a strict intersection logic that encodes occurrence within the sampled duplicate classes across a fixed set of taxa.

For a fixed set of taxa:

- For each *Arabidopsis* identifier, construct a binary presence-absence vector.
- Group identifiers with identical vectors.
- Each group defines a strict intersection occupancy class.

This procedure partitions the distinct Arabidopsis identifiers from the sampled duplication-mode sets into equivalence classes defined solely by shared taxonomic presence patterns. These classes form the mutually exclusive units from which the downstream functional foregrounds were assembled.

Properties:

- Each gene belongs to exactly one occupancy class.
- Occupancy classes are mutually exclusive.
- These classes are never modified downstream.

This binning step fixes mutually exclusive intersections of gene-identifier occupancy across taxa; it does not preclude later intersections of independently FDR-significant GO term sets. By fixing occupancy class membership at this stage, all subsequent analyses inherit a stable, reproducible partitioning of genes that is independent of downstream biological hypotheses.

Later GO-term intersections summarize terms independently significant in multiple tested taxa; they do not modify the gene-level occupancy classes.

#### Foreground Construction as Unions of Occupancy Classes

Genome-derived GO foregrounds were formed by deterministic unions of exact nine-taxon duplicate-mode occupancy classes. A gene may contribute to more than one higher-level species foreground, but each distinct Arabidopsis homolog is counted only once within a given foreground. GO enrichment is computed separately for each taxon and duplication mode.

Explicitly:

- No genes were intersected across occupancy classes.
- No classes were intersected with other classes.
- Foregrounds are deterministic unions defined by occupancy class membership.

The exact occupancy-class representation prevents double counting of Arabidopsis homolog identifiers within a defined foreground. Absence from a duplication-mode-specific class does not establish whole-genome absence.

The same exact-class lookup is used for syntenic and tandem gene sets before standardized GO testing.

- duplication-mode comparisons
- local retention classifications, used descriptively and not as phased subgenome categories
- lineage-associated duplicate-mode gene sets

Different biological comparisons are interpreted at the level of their explicit gene sets; statistical enrichment is not claimed for untested subgenome-defined gene categories.

#### GO Enrichment

Genome-derived GO enrichment was calculated using the fixed Arabidopsis GO annotation file dated 1 October 2025 and go-basic ontology dated 10 October 2025. Only direct, non-negated Biological Process associations to active GO terms were included, without ancestor propagation. For each of five taxa (*Sarracenia*, *Nepenthes*, *Pinguicula*, *Utricularia* and *Actinidia*) and each duplication mode, the exact-class-union homolog foreground was restricted to 23,167 eligible Arabidopsis genes with Biological Process annotations.

For each of 3,708 Biological Process terms represented in the background, a one-sided hypergeometric overrepresentation P value was calculated independently for each taxon and duplication mode. Terms absent from a foreground were retained with P = 1.

- The foreground hit count k is the number of distinct Arabidopsis homologs annotated directly to the term.
- The background hit count K is the number of the 23,167 annotated background homologs assigned directly to the term.
- The hypergeometric survival probability P(X >= k) used the foreground size n and the fixed annotated background size N = 23,167.

The same 23,167-gene, 3,708-term annotated background was used for all ten taxon/duplication-mode testing families. It is an explicitly standardized Arabidopsis background to ensure identical testing denominators across the five species.

Benjamini-Hochberg false discovery rate correction was performed separately over all 3,708 tests within each taxon and duplication mode. A category was considered overrepresented when BH q < 0.05.

Expression-associated GO enrichment is a separate analytical branch and is not inferred from the nine-taxon duplicate-mode tests. The standardized rarefaction analysis described below also uses GO annotations but measures descriptive annotation breadth rather than enrichment.

A *Sarracenia*-*Actinidia* comparison was tabulated for each duplication mode after the independent within-taxon tests. Terms with *Sarracenia* q < 0.05 but *Actinidia* q >= 0.05 were retained as *Sarracenia*-enriched, *Actinidia*-nonsignificant categories; this is not interpreted as biological absence from *Actinidia*.

### RNA-seq Processing and Expression Analyses

#### RNA Sequencing and Sample Design

RNA sequencing was performed on three tissue types: roots, trap leaves, and flowers. Three biological replicates were collected for each tissue type, yielding a total of nine RNA-seq samples. Samples were designated as follows: SPR1-SPR3 (roots), SPL1-SPL3 (trap leaves), and SPF1-SPF3 (flowers). Flowers were sampled after petal abscission, representing the post-anthesis stage.

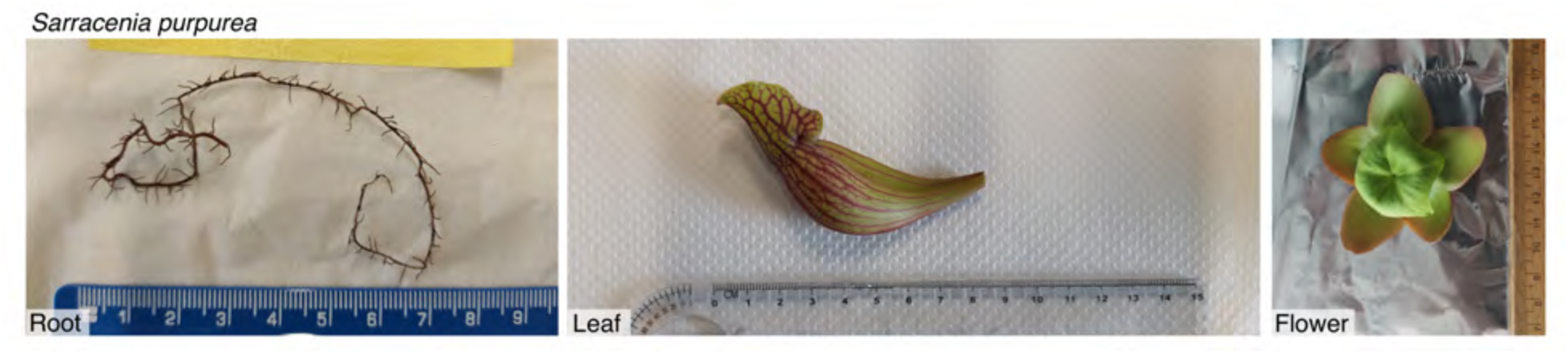

RNA-seq libraries were sequenced on an Illumina platform using paired-end reads. All RNA-seq data have been deposited under BioProject PRJDB15737. Detailed accession information for each sample is provided in section 4.9.

#### RNA-seq Samples and Accession Metadata

The nine RNA-seq libraries and their BioSample, experiment, run, and FASTQ identifiers are listed here:

| **TISSUE** | **SAMPLE** | **BIOSAMPLE** | **EXPERIMENT** | **RUN** | **LIBRARY LAYOUT** | **READ FILES** |
|---|---|---|---|---|---|---|
| **FLOWER** | SPF1 | SAMD00754511 | DRX522175 | DRR538356 | Paired-end | SPF1_1.fq.gz, SPF1_2.fq.gz |
| **FLOWER** | SPF2 | SAMD00754512 | DRX522176 | DRR538357 | Paired-end | SPF2_1.fq.gz, SPF2_2.fq.gz |
| **FLOWER** | SPF3 | SAMD00754513 | DRX522177 | DRR538358 | Paired-end | SPF4_1.fq.gz, SPF4_2.fq.gz |
| **LEAF** | SPL1 | SAMD00754514 | DRX522178 | DRR538359 | Paired-end | SPL1_1.fq.gz, SPL1_2.fq.gz |
| **LEAF** | SPL2 | SAMD00754515 | DRX522179 | DRR538360 | Paired-end | SPL2_1.fq.gz, SPL2_2.fq.gz |
| **LEAF** | SPL3 | SAMD00754516 | DRX522180 | DRR538361 | Paired-end | SPL3_1.fq.gz, SPL3_2.fq.gz |
| **ROOT** | SPR1 | SAMD00754517 | DRX522181 | DRR538362 | Paired-end | SPR1_1.fq.gz, SPR1_2.fq.gz |
| **ROOT** | SPR2 | SAMD00754518 | DRX522182 | DRR538363 | Paired-end | SPR2_1.fq.gz, SPR2_2.fq.gz |
| **ROOT** | SPR3 | SAMD00754519 | DRX522183 | DRR538364 | Paired-end | SPR3_1.fq.gz, SPR3_2.fq.gz |

#### RNA-seq Read Mapping and Transcriptome Assembly

Genome indexing for RNA-seq read alignment was performed using HISAT2 v2.2.1, incorporating splice-site and exon information derived from the final EVM-integrated gene annotation. Splice sites were extracted from the GFF3 annotation using *hisat2_extract_splice_sites.py*. Exon coordinates were obtained by converting the GFF3 file to GTF format using gffread v0.12.7, followed by *hisat2_extract_exons.py*.

The RagTag-scaffolded genome assembly was indexed using:

hisat2-build --ss Spur.ss --exon Spur.exons Spur.fasta --large-index -p 16 Spur_index RNA-seq reads were aligned to the indexed genome using HISAT2 with parameters optimized for transcriptome assembly:

hisat2 --dta -p 16 -x Spur_index \

-1 SAMPLE_R1.fastq.gz -2 SAMPLE_R2.fastq.gz \

-S SAMPLE.sam

The --dta option was used to report alignments suitable for transcript assemblers. SAM files were converted to BAM format and sorted using SAMtools v1.16.1.

Transcript assembly and expression estimation were performed independently for each sample using StringTie v2.2.1: stringtie SAMPLE.sorted.bam -G reference.gtf -e -B -o SAMPLE.gtf

The -e option restricted assembly to reference transcripts, and the -B option generated tabular output files required for downstream differential expression analyses.

#### Differential Expression Analysis

Gene- and transcript-level count matrices were generated using *prepDE.py* following the StringTie workflow. The gene-level count matrix was analyzed using DESeq2 v1.36.0 in R v4.2.0 (Love et al., 2014), with three biological replicates each from flower, pitcher leaf, and root tissues. The original flower sample SPF4 was relabeled SPF3 for analysis, explaining the SPF4-named read files associated with sample SPF3 in the accession table. The annotation contains one representative transcript per gene; therefore, differential expression is reported at the gene level.

Of 28,835 genes in the input count matrix, 22,910 were retained after excluding genes with fewer than 10 counts summed across the nine samples. Sample-specific size factors were estimated by DESeq2’s default median-of-ratios normalization method. Counts were modeled using a negative binomial generalized linear model with tissue as the sole explanatory variable (design = ∼ tissue). Flower-versus-pitcher-leaf, root-versus-pitcher-leaf, and flower-versus-root contrasts were evaluated using Wald tests. Default independent filtering by mean normalized count was enabled, and P values were adjusted independently within each contrast using the Benjamini– Hochberg method. Genes with adjusted P < 0.05 in at least one contrast formed the combined 17,729-gene DEG table. Thirty-six of these genes lacked adjusted P values for root versus pitcher leaf because of independent filtering but met the significance criterion in at least one other contrast.

For the flower-versus-pitcher-leaf volcano plot (**Figure 5**), colored upregulation classifications additionally required |log₂ fold change| > 1. For the duplicate-class expression-responsiveness analysis (**Figure 6A**), genes significant in at least one contrast were intersected with the syntenic and tandem duplicate classes; responsiveness was calculated as the maximum absolute log₂ fold change among the three contrasts. For **Supplementary Figure S7**, log₂ fold changes for the selected 50 most flower-biased and 50 most pitcher-leaf-biased genes were standardized as z scores across the 100 genes separately within each pairwise comparison.

For exploratory principal component analysis and sample-distance heatmaps, expression values were variance-stabilized with DESeq2’s vst function (blind = FALSE), and Euclidean sample distances were computed from those transformed profiles. Where variance-stabilized gene- expression heatmaps were constructed with pheatmap, values were standardized within each gene across samples; this approach is distinct from the comparison-wise log₂-fold-change standardization used for **Supplementary Figure S7**. All reported DESeq2 log₂ fold changes were unshrunken; no lfcShrink() was applied.

#### Expression Responsiveness of Syntenic Versus Tandem Duplicates

The primary comparison used the gene-level dataset

*Sarracenia*_syntenic_vs_tandem_expression_table.tsv, which records the original tandem and syntenic classifications and expression responsiveness for 934 and 4,578 *Sarracenia* genes, respectively. This dataset is available with the reproducibility code. All included genes have adjusted P < 0.05 for at least one of the three tissue contrasts reported in all_DEGs.tsv.

For each selected gene, responsiveness was computed as max(|FvL_log2FC|, |RvL_log2FC|, |FvR_log2FC|) from all_DEGs.tsv, representing flower versus leaf, root versus leaf, and flower versus root, respectively. All 5,512 records match this definition exactly. No additional pseudocount was applied; expression values were not recalculated from the underlying read-count matrix. This statistic measures the largest pairwise tissue-associated fold change, not expression direction or a change unique to pitchers.

The duplication classes were compared using gene-level maximum absolute log2 fold changes. A two-sided Mann-Whitney U test evaluated class differences, and Cliff’s delta was calculated as 2U/(n_tandem × n_syntenic) - 1, using mean ranks for ties. The test describes differences in the selected gene population; genes sharing duplication history or close homology need not be evolutionarily independent.

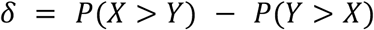

The sample contains 934 tandem genes and 4,578 syntenic genes, with no subsampling for the expression test. Their values yield Mann-Whitney P = 6.937157 × 10^-18 and Cliff’s delta = 0.178618 (**Figure 6A**).

#### Functional Breadth Analysis Using GO Rarefaction

For the historical rarefaction we mapped the expression-responsive *Sarracenia* gene IDs to one-best-hit Arabidopsis homolog IDs using *Sarracenia*_to_Araport11.txt, derived from the archived filtered *Sarracenia*-Araport11 protein BLAST results. Multiple *Sarracenia* genes mapping to a single Arabidopsis locus were collapsed to one homolog per duplication-class set. Direct Biological Process (aspect P) assignments were extracted from the TAIR gene-association GAF generated 1 October 2025. The historical script did not explicitly exclude negated annotations or obsolete identifiers, unlike the separately standardized GO overrepresentation pipeline. After mapping and GO annotation, the sampling pools comprised 617 tandem-associated and 2,305 syntenic-associated Arabidopsis homologs.

For each class and sampling depth, we drew 300 random samples without replacement; the archived script uses seed 2 for tandem and seed 1 for syntenic samples. Richness was the number of distinct observed Biological Process terms. Effective breadth D1 was exp(H), where H is Shannon entropy calculated from each term’s relative proportion of gene-by-term annotation incidences (Jost 2006; Chao, et al. 2014). Rarefaction at equal numbers of annotated Arabidopsis homologs controls the number of sampled homolog IDs, not the number of *Sarracenia* paralogs.

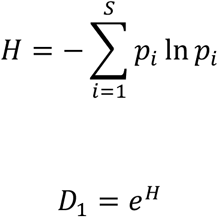

For each rarefaction replicate, each observed homolog-by-GO-term incidence contributes once to the term-frequency distribution. D1 therefore represents the effective number of annotated GO categories under this mapping and ontology snapshot, rather than a direct measurement of ecological functional diversity.

For each class and sampling depth, we plot the archived mean and 2.5th-97.5th percentiles of the 300 subsamples. Rerunning the code and inputs reproduces the directions and approximate magnitudes of the archived curves, with small differences in stochastic means. Subsamples from one underlying gene set are not independent biological replicates; accordingly, we do not use a Mann-Whitney test between the replicate subsamples to infer a population-level difference in rarefaction curves. This descriptive analysis is restricted to the expression-responsive duplicate-class gene sets.

#### Use of Large Language Model Tools

OpenAI ChatGPT (web application; accessed during manuscript preparation and revision in 2026) was used interactively to assist with development and checking of bioinformatic approaches and analysis scripts, preparation of selected graphics, and editorial review of manuscript text. Queries were natural-language prompts specifying the biological or analytical objective, relevant input files, requested outputs, statistical or formatting constraints, and, where appropriate, requests for verification against underlying data. Generated scripts, analyses, graphics, and prose were reviewed, tested, and edited by the authors before inclusion. Earlier exploratory ChatGPT interactions used the then-current models available through the service; no model was treated as a source of scientific evidence. The authors take responsibility for all analyses, interpretations, and text presented here.

### Data availability

The *Sarracenia purpurea* genome assembly and annotation are available from CoGe (https://genomevolution.org/coge/GenomeInfo.pl?gid=69349) and the European Nucleotide Archive (ENA) Research Project Accession PRJEB128613 and ERP208013. The tissue sample accession is ERS32160899 and sequencing run accession is ERR18068765. The analysis accession is ERZ30649059 (the annotated assembly). The RNA-seq data are associated with BioProject PRJDB15737. Reproduction code, analytical inputs, RepeatMasker GFF/.out/.tbl outputs, SynMap and FractBias exports, GO reference snapshots, gene-level DEG results, and detailed RNA-seq workflow pseudocode are available in Zenodo (DOI 10.5281/zenodo.24475066.) The archived README identifies the executable workflows and public upstream accessions.

## Acknowledgments

We acknowledge the following sources for funding: the U.S. National Science Foundation grants 2030871 and 2139311 (to VAA and CL, respectively), the Sofja Kovalevskaja Program of the Alexander von Humboldt Foundation (to KF), and JSPS KAKENHI 23K20050 (to KF).

## Author contributions

VAA conceived the study; VAA led the research with assistance from JK; VAA, JK, MR, and CL provided materials; MR sequenced Oxford Nanopore reads; KF and MF provided RNA-seq data; CP, JK, and NP assembled and annotated the genome; CP decontaminated and reference- scaffolded the assembly; CP analyzed RNA-seq data; VAA, JK, CP, NP, and JM performed additional downstream bioinformatic analyses; VAA, CP, and JM produced graphics; VAA wrote the manuscript, and all authors approved the final version.

## Conflict of interest

The authors declare no competing interests.

## Supplementary material for

### Supplementary material

**Supplementary Figure S1.**
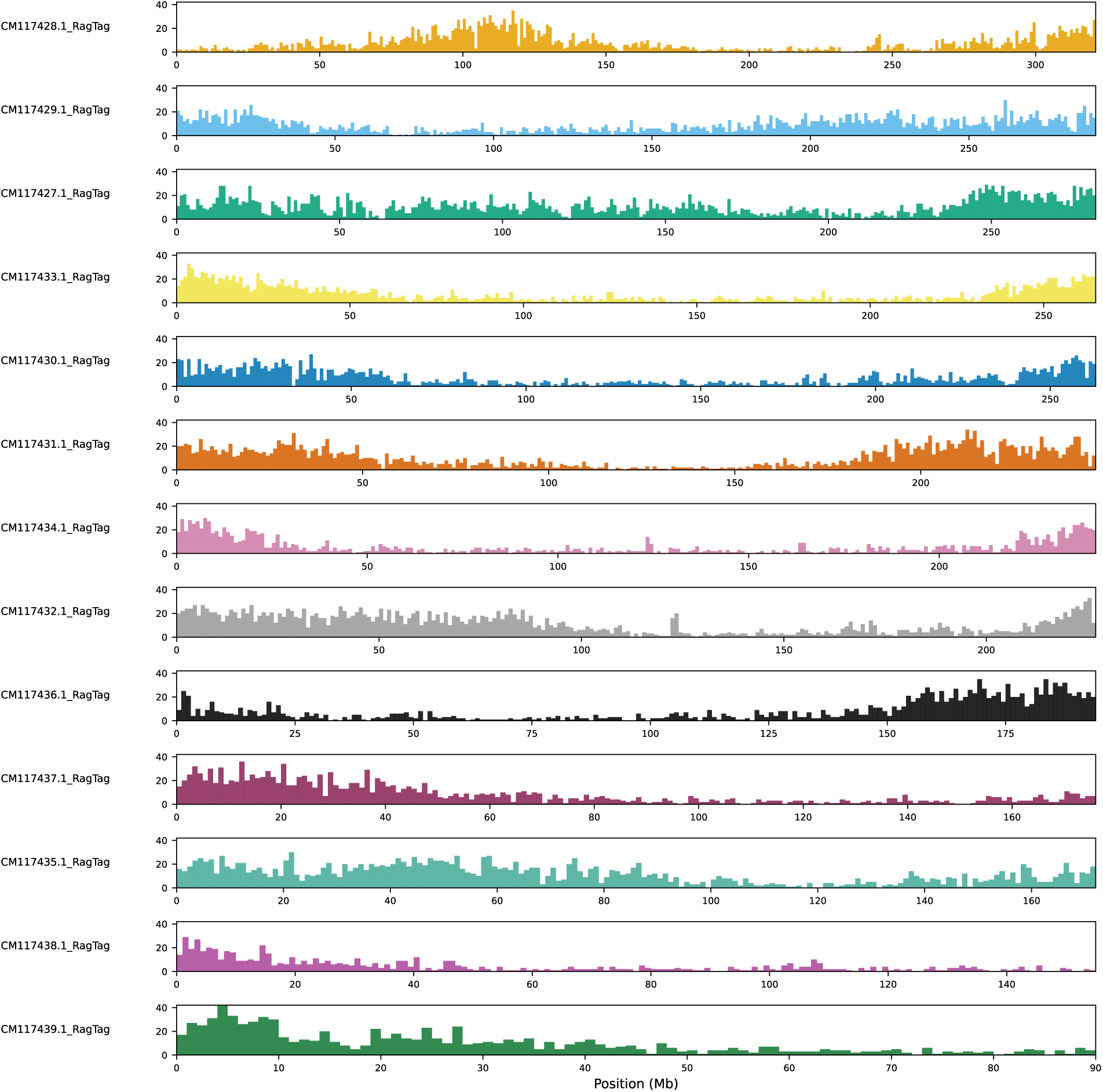
Gene-density variation across the 13 primary *Sarracenia purpurea* pseudochromosomes. Gene counts per 1-Mb window across the 13 pseudochromosomes highlight extended gene-poor regions and localized peaks of elevated density.

**Supplementary Figure S2.**
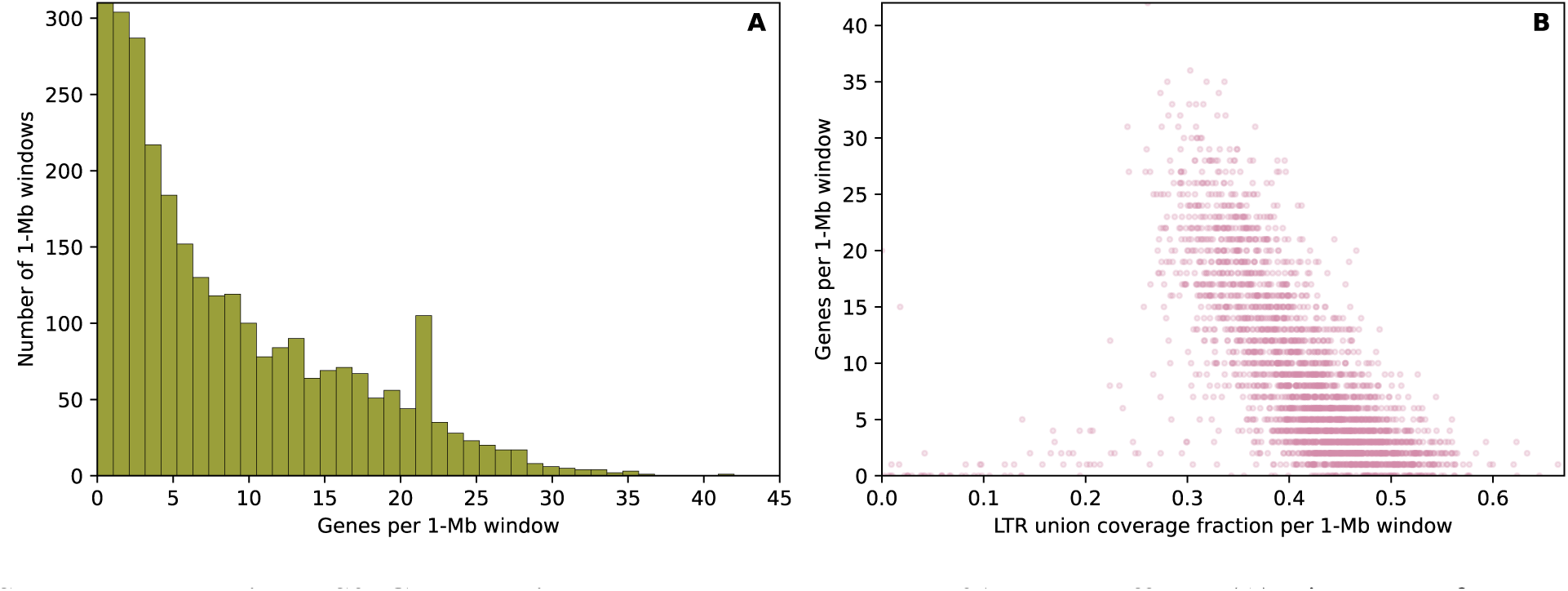
Gene density and LTR coverage across 31 large scaffolds. (**A**) Histogram of genes overlapping 1-Mb windows across 31 scaffolds longer than 5 Mb (3,211 windows). (**B**) Gene count versus the nonoverlapping fraction of each window covered by RepeatMasker fragments explicitly tagged as LTR (Spearman ρ = −0.671). The LTR-tagged subset is narrower than the full LTR-retrotransposon class.

**Supplementary Figure S3.**
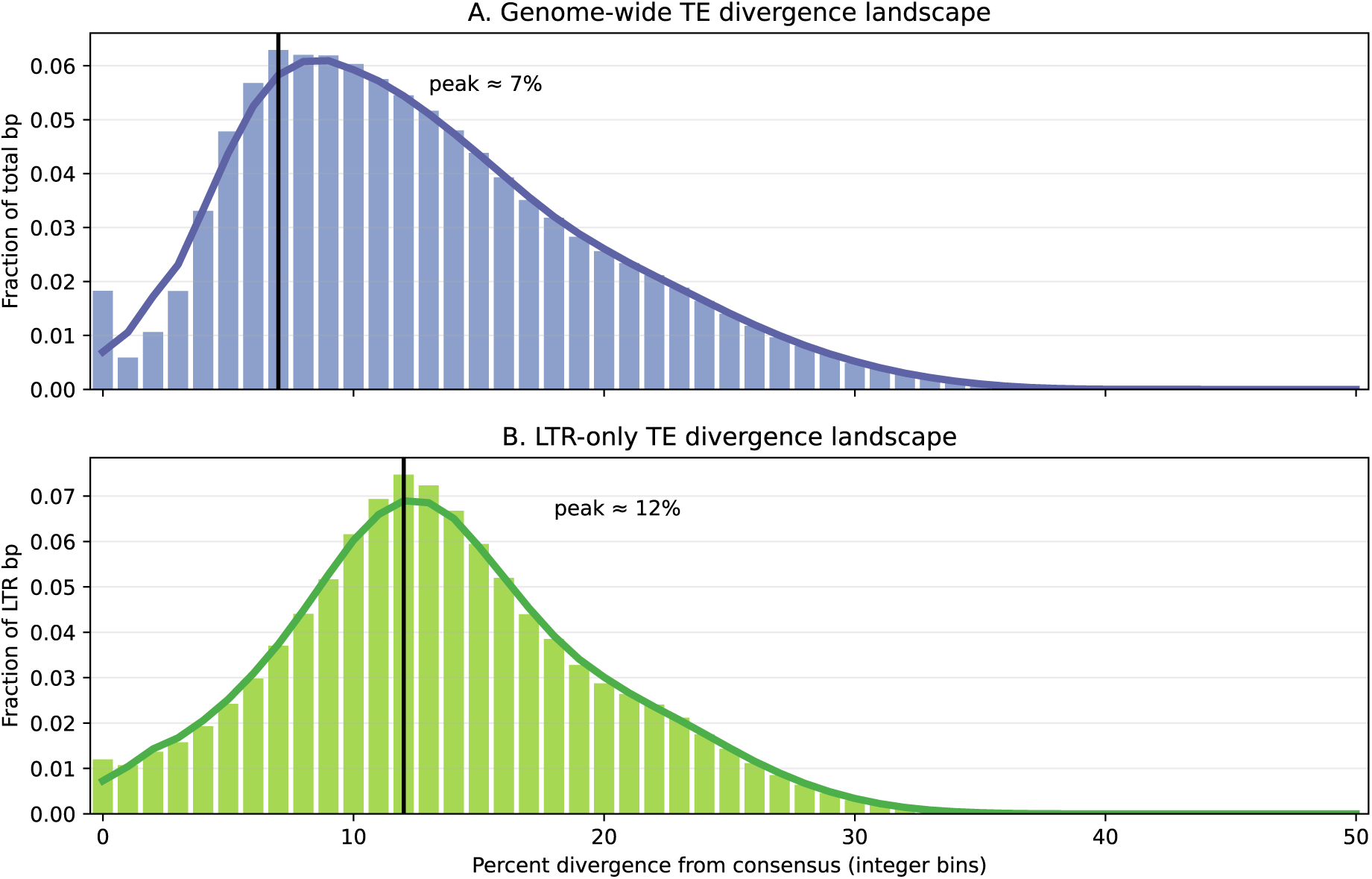
Repeat divergence landscapes for *Sarracenia purpurea* repeats (genome-wide versus LTR-tagged). (A) Genome-wide and (B) LTR-tagged bp-weighted divergence distributions derived from RepeatMasker annotations; vertical lines mark modal divergence bins. For each RepeatMasker-annotated repeat fragment, the reported percent divergence from its consensus was binned into integer divergence bins (0-50%), and base-pair coverage was summed per bin to generate bp-weighted divergence profiles. Histograms show the fraction of total covered bp contributed by each divergence bin (panel-specific normalization), and the overlaid smoothed curves summarize the same distributions to highlight modal structure. Vertical reference lines mark the peak (modal) divergence bin for each panel. The LTR-tagged subset consists of RepeatMasker fragments whose repeat names explicitly contain “LTR” and is narrower than the complete LTR-retrotransposon class reported in Table S2.

**Supplementary Figure S4.**
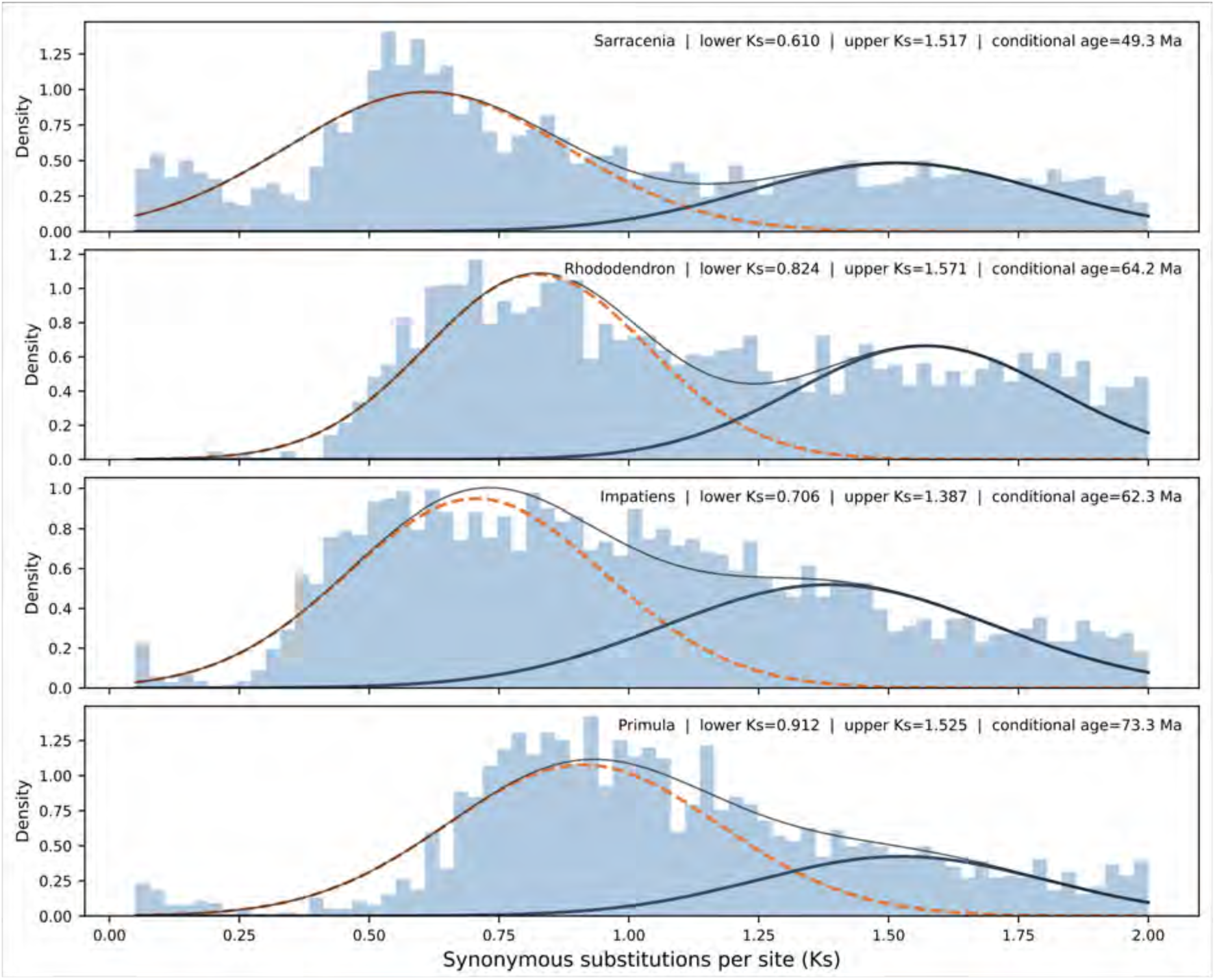
Self-synteny Ks distributions across four selected Ericales taxa. Self-syntenic gene-pair Ks values were restricted to 0.05 ≤ Ks ≤ 2.0 before fitting two-component Gaussian mixture models. Blue histograms show the observed Ks distributions; dashed orange and solid dark curves represent the lower- and upper-Ks components, respectively, with the gray curve showing their sum. Conditional younger-component ages for *Sarracenia* (49.3 Ma), *Rhododendron* (64.2 Ma), *Impatiens* (62.3 Ma), and *Primula* (73.3 Ma) assume that the respective upper-Ks components correspond to the core-eudicot *gamma* hexaploidy event at 122.5 Ma. Component homology and inferred absolute ages are model- and calibration-dependent; two components were selected for biological interpretation and do not necessarily minimize the Bayesian information criterion (BIC).

**Supplementary Figure S5.**
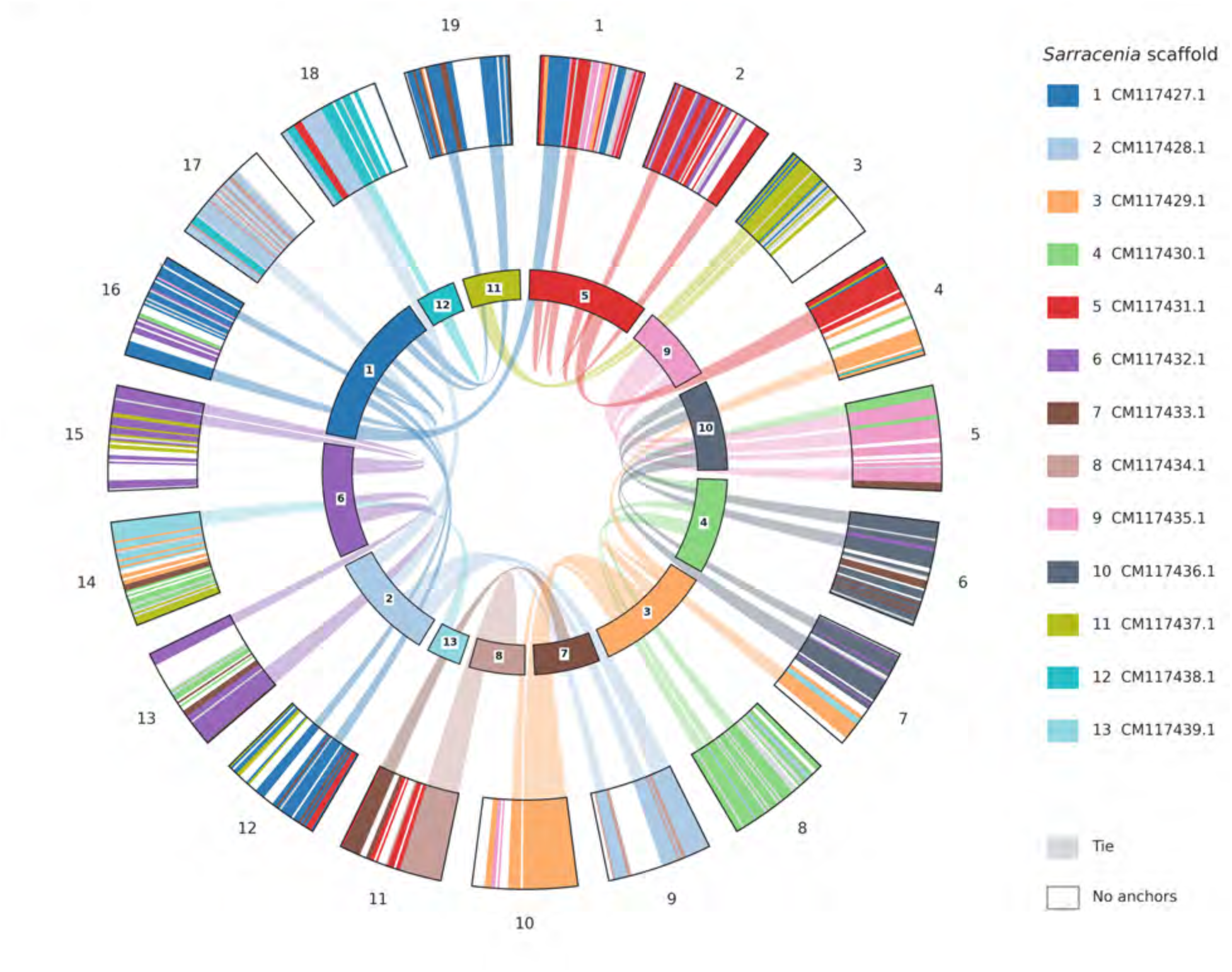
Locally highest-retention *Sarracenia* scaffold along the *Vitis* reference. Each of the 19 Vitis chromosomes was partitioned into 50 physical-coordinate bins using CoGe genome 19990. Colors distinguish the 13 *Sarracenia* primary scaffolds with the uniquely highest finite-Ks syntenic-anchor count in each bin; gray denotes a tied highest count and white denotes no qualifying anchor. Changes in the leading scaffold show locally heterogeneous retention and rearrangement, not phased ancestral dominant and recessive parental subgenomes. The inner sectors represent the 13 *Sarracenia* scaffolds, and 42 ribbons join runs of at least five consecutive uniquely assigned bins, spanning 340 bins; the five-bin cutoff affects ribbons only. Of 950 reference bins, 688 have a unique leading scaffold, 55 are tied, and 207 lack qualifying anchors.

**Supplementary Figure S6.**
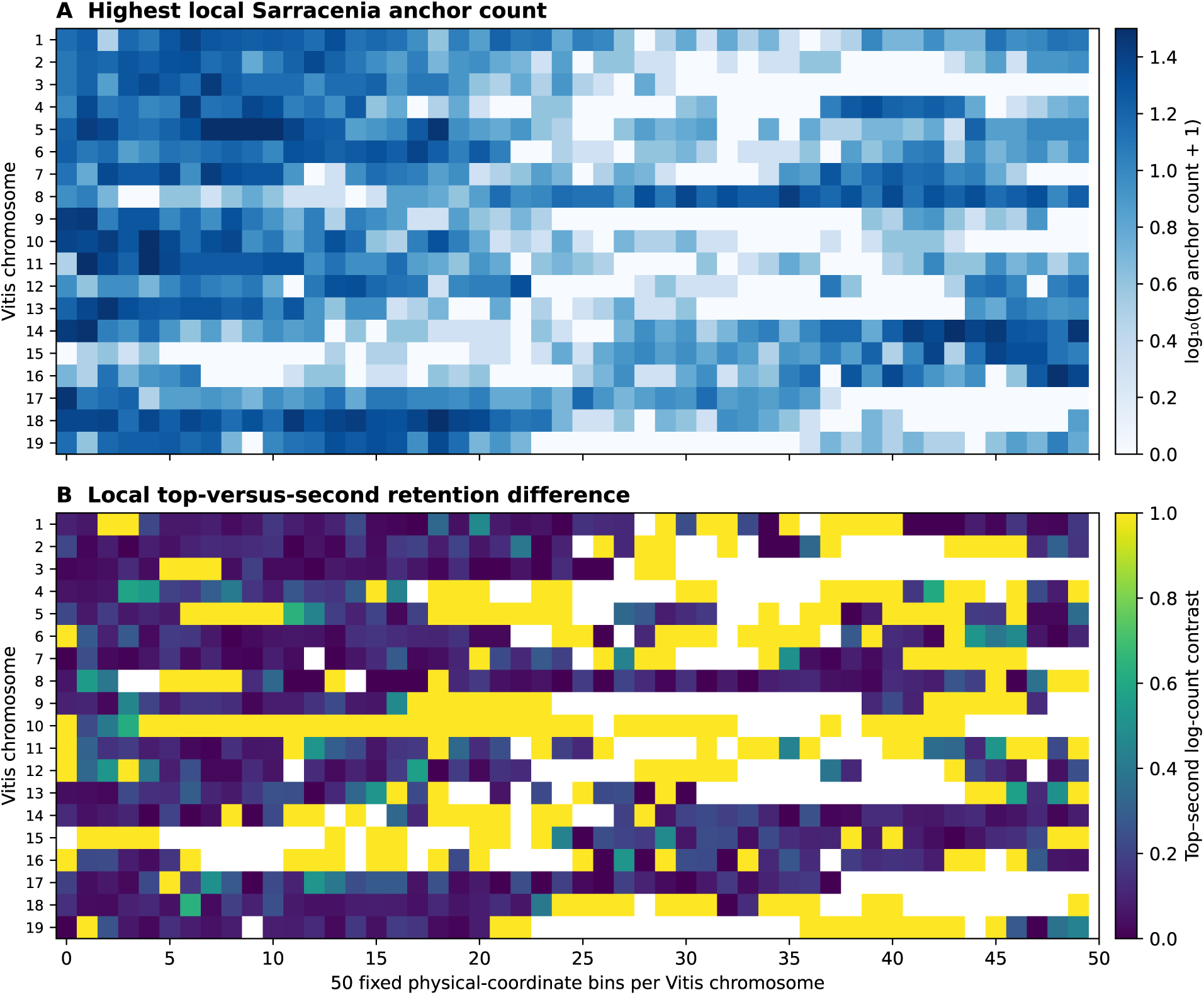
*Sarracenia*–*Vitis* syntenic-retention heatmaps under the manuscript-aligned 50-bin calculation. (A) Log10-transformed anchor counts for the locally highest-retention *Sarracenia* pseudochromosome within each *Vitis* reference-coordinate bin. (B) Difference between the largest and second-largest log10(anchor count + 1) values, normalized by their sum; white cells lack qualifying finite-Ks anchors. The contrast is sensitive to bins represented by only one positive *Sarracenia* track; these data do not identify progenitor-derived dominant or recessive subgenomes.

**Supplementary Figure S7.**
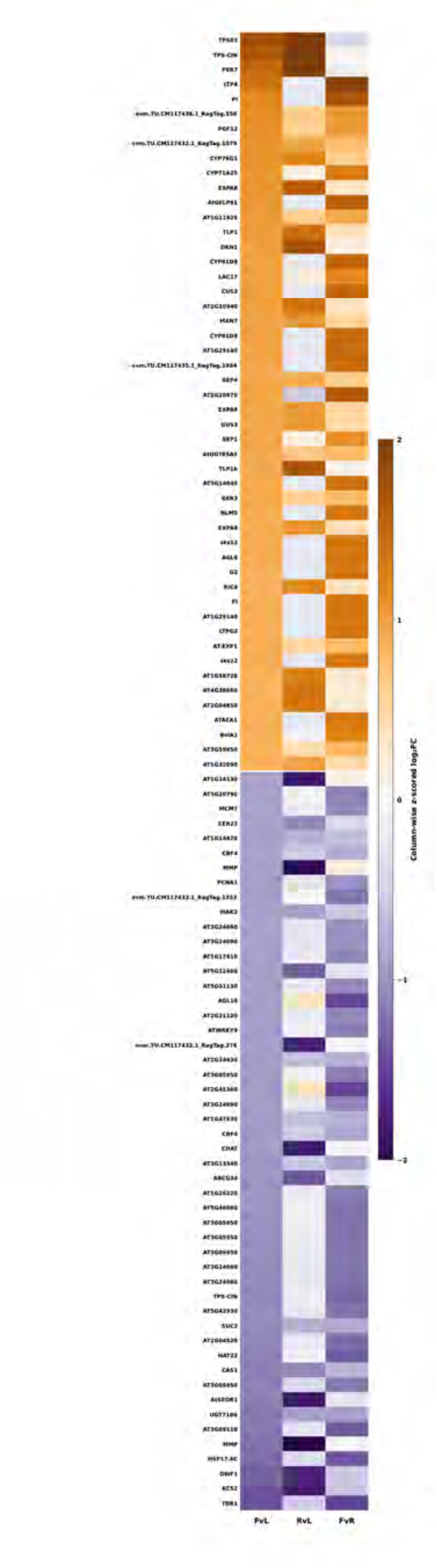
Differential-expression patterns across flowers, roots, and pitcher leaves in *Sarracenia purpurea*. Heatmap showing 100 genes selected according to the magnitude of their flower-versus-pitcher-leaf (FvL) expression differences. The 50 most flower-biased genes are ordered from highest to lowest positive log₂ fold change (log₂FC), followed by the 50 most pitcher-biased genes, ordered from less negative to more negative FvL values. All selected genes have FvL-adjusted P < 0.05. Columns represent flowers versus pitcher leaves (FvL), roots versus pitcher leaves (RvL), and flowers versus roots (FvR). Within each comparison, log₂FC values were independently standardized across the 100 selected genes by subtracting the column mean and dividing by its standard deviation. The color scale ranges from −2 to +2, with values beyond these limits clipped for visualization. Orange and purple indicate relatively higher and lower standardized values, respectively, rather than necessarily positive and negative raw fold changes. Gene labels correspond to best Arabidopsis sequence matches, with *Sarracenia* identifiers retained when no match was available. Repeated Arabidopsis annotations can represent distinct *Sarracenia* genes. Flower-biased genes include homologs annotated as *PI*, *SEP1*, *SEP4*, *TPS03*, *TPS-CIN*, *PER7*, and *LTP4*, whereas strongly pitcher-biased genes include homologs associated with stress responses, metabolism, and signaling.

### Supplementary Tables

**Table S1.** Summary of gene and transcript structural features in the *Sarracenia purpurea* genome. Key annotation metrics for the primary gene set, including counts and length distributions for genes, transcripts, coding sequences, exons, and introns. Values summarize overall genome architecture, highlighting a moderately exon-rich gene space, substantial intron-length variation, and a genome-wide gene density of about 8 genes per megabase across the 3.52 Gb assembly. Each gene locus was represented by a single representative transcript model in the final annotation.

| **METRIC** | **VALUE** |
|---|---|
| **NUMBER OF GENE LOCI** | 28,835 |
| **NUMBER OF TRANSCRIPTS** | 28,835 |
| **TOTAL CDS FEATURES** | 132,001 |
| **MEAN EXON LENGTH (BP)** | 263.12 |
| **MEDIAN EXON LENGTH (BP)** | 139 |
| **MEAN GENE LENGTH (BP)** | 9,315.68 |
| **MEDIAN GENE LENGTH (BP)** | 3,112 |
| **MEAN TRANSCRIPT LENGTH (BP)** | 9,315.68 |
| **MEDIAN TRANSCRIPT LENGTH (BP)** | 3,112 |
| **MEAN EXON COUNT PER GENE** | 4.58 |
| **MEDIAN EXON COUNT PER GENE** | 3 |
| **NUMBER OF SINGLE-EXON GENES** | 7,179 |
| **MEAN INTRON LENGTH (BP)** | 2,267.08 |
| **MEDIAN INTRON LENGTH (BP)** | 558 |
| **GENOME-WIDE GENE DENSITY (GENES/MB)** | 8.18 |
| **TOTAL ASSEMBLY LENGTH (BP)** | 3,523,999,544 |

**Table S2.** Genome-wide distribution of transposable element classes and other masked repeat categories in the *Sarracenia purpurea* assembly. Summary of RepeatMasker annotations showing the proportional contributions of major TE groups, including LTR retrotransposons, LINEs, SINEs, DNA transposons, rolling-circle elements, and unclassified repeats. The table also reports ancillary repeat categories such as small RNAs, simple repeats, and low-complexity regions, providing an overview of the repeat landscape that shapes genome size, structure, and gene-density variation in *Sarracenia purpurea*. Repeat-class counts and masked-base percentages were obtained from the original RepeatMasker v4.1.5 .tbl summary, using 3,523,999,544 bp across 2,042 input sequences as the reference length; the final scaffolded assembly has 3,523,623,004 bp across 1,884 scaffolds. The original .out, .tbl, and GFF outputs accompany the reproduction archive.

| **REPEAT CATEGORY** | **NUMBER OF ELEMENTS** | **BASES OCCUPIED (BP)** | **% OF GENOME** |
|---|---|---|---|
| **TOTAL MASKED SEQUENCE** | - | 3,099,995,857 | 87.97 |
| **TOTAL INTERSPERSED REPEATS** | - | 3,051,425,825 | 86.59 |
| **RETROELEMENTS (TOTAL)** | 3,018,215 | 2,578,482,566 | 73.17 |
| **SINES** | 8,757 | 9,815,940 | 0.28 |
| **LINES** | 63,405 | 38,409,777 | 1.09 |
| **LTR RETROTRANSPOSONS (TOTAL)** | 2,946,053 | 2,530,256,849 | 71.80 |
| **TY1/COPIA** | 1,143,498 | 1,176,556,747 | 33.39 |
| **GYPSY/DIRS1** | 555,662 | 721,463,009 | 20.47 |
| **RETROVIRAL** | 16,784 | 10,010,854 | 0.28 |
| **DNA TRANSPOSONS (TOTAL)** | 139,026 | 36,995,745 | 1.05 |
| **HOBO-ACTIVATOR** | 20,275 | 11,713,441 | 0.33 |
| **MULE-MUDR** | 9,274 | 6,885,918 | 0.20 |
| **OTHER DNA TRANSPOSONS** | - | 18,396,386 | 0.52 |
| **ROLLING-CIRCLE ELEMENTS** | 7,995 | 2,565,999 | 0.07 |
| **UNCLASSIFIED INTERSPERSED REPEATS** | 1,508,633 | 435,947,514 | 12.37 |
| **SMALL RNA** | 11,922 | 25,821,641 | 0.73 |
| **SIMPLE REPEATS** | 481,899 | 21,756,034 | 0.62 |
| **LOW-COMPLEXITY SEQUENCE** | 53,438 | 2,867,746 | 0.08 |
| **SATELLITES** | 264 | 79,125 | <0.01 |

**Table S3.** GO Biological Process terms enriched in *Sarracenia* duplicate-class homologs but not significantly enriched in the corresponding Actinidia comparison. Genome-derived GO terms were tested independently using the frozen 2025 TAIR/GO framework and BH adjustment over 3,708 terms in each taxon and duplication mode. Each row represents a GO-term/duplication-mode association; Actinidia non-significance does not imply genomic or biological absence. This table replaces the historical six-category summary, which was not reproduced under the standardized framework.

| **DUPLICATION MODE** | **GO IDENTIFIER** | **BIOLOGICAL PROCESS** | ***SARRACENIA* BH Q** | ***ACTINIDIA* BH Q** |
|---|---|---|---|---|
| **SYNTENIC** | GO:0000122 | negative regulation of transcription by RNA polymerase II | 0.03698 | 0.3783 |
| **SYNTENIC** | GO:0009789 | positive regulation of abscisic acid-activated signaling pathway | 0.0001696 | 0.07308 |
| **SYNTENIC** | GO:0009863 | salicylic acid mediated signaling pathway | 0.02489 | 0.06446 |
| **SYNTENIC** | GO:0010073 | meristem maintenance | 0.0175 | 0.05194 |
| **SYNTENIC** | GO:0010199 | organ boundary specification between lateral organs and the meristem | 0.01848 | 0.221 |
| **SYNTENIC** | GO:0010214 | seed coat development | 0.0256 | 0.1064 |
| **SYNTENIC** | GO:0010321 | regulation of vegetative phase change | 0.03303 | 0.6705 |
| **SYNTENIC** | GO:0010508 | positive regulation of autophagy | 0.03303 | 0.2291 |
| **SYNTENIC** | GO:0010588 | cotyledon vascular tissue pattern formation | 0.01795 | 0.06446 |
| **SYNTENIC** | GO:0030308 | negative regulation of cell growth | 0.01848 | 0.221 |
| **SYNTENIC** | GO:0042335 | cuticle development | 0.009586 | 0.1322 |
| **SYNTENIC** | GO:0048467 | gynoecium development | 0.03313 | 0.2291 |
| **SYNTENIC** | GO:0048765 | root hair cell differentiation | 0.009586 | 0.2291 |
| **SYNTENIC** | GO:0051512 | positive regulation of unidimensional cell growth | 0.0149 | 0.1306 |
| **SYNTENIC** | GO:0060586 | multicellular organismal-level iron ion homeostasis | 0.03163 | 0.08465 |
| **SYNTENIC** | GO:0071244 | cellular response to carbon dioxide | 0.03303 | 0.06966 |
| **SYNTENIC** | GO:0080156 | mitochondrial mRNA modification | 0.0002897 | 0.3783 |
| **SYNTENIC** | GO:1901002 | positive regulation of response to salt stress | 0.02879 | 0.5921 |
| **TANDEM** | GO:0002679 | respiratory burst involved in defense response | 0.03372 | 0.7022 |
| **TANDEM** | GO:0006749 | glutathione metabolic process | 0.03372 | 0.8321 |
| **TANDEM** | GO:0006857 | oligopeptide transport | 0.03372 | 1 |
| **TANDEM** | GO:0006952 | defense response | 0.03372 | 0.06089 |
| **TANDEM** | GO:0006979 | response to oxidative stress | 0.01702 | 0.3641 |
| **TANDEM** | GO:0009620 | response to fungus | 0.0411 | 0.08785 |
| **TANDEM** | GO:0009695 | jasmonic acid biosynthetic process | 0.03372 | 0.129 |
| **TANDEM** | GO:0010150 | leaf senescence | 0.001592 | 0.5256 |
| **TANDEM** | GO:0016554 | cytidine to uridine editing | 0.004388 | 1 |

**Supplementary Tables S4–S12** are provided as individually named CSV files, Supplementary_Table_S4.csv through Supplementary_Table_S12.csv.

## Notes

### Competing Interest Statement

The authors have declared no competing interest.

### Summary of Updates

Main manuscript updated and merged with supplement

